# Differential regulation of KCC2 function, trafficking, and degradation by Ca^2+^-dependent signaling pathways

**DOI:** 10.64898/2026.07.02.736223

**Authors:** Marc J. Bergeron, Isabel Plasencia-Fernandez, Annie Barbeau, Noémie Comeau, Martin Cottet, Antoine G. Godin, Yves De Koninck

## Abstract

Regulation of the K^+^-Cl^-^ cotransporter KCC2 is a critical determinant of the efficacy of inhibition in the central nervous system and KCC2 hypofunction appears at the root of several neurological disorders. Both BDNF-TrkB and NMDAR signaling regulate KCC2, but how they interact remains unknown. Here we show that these two signaling pathways act synergistically to differentially modulate KCC2 function and expression through post-translational regulation, via distinct Ca^2+^ signalling modes. Blocking ryanodine-dependent intracellular Ca^2+^ release prevented TrkB-, but not NMDAR-mediated downregulation. TrkB-signalling in absence of NMDAR activation modulated KCC2 function but not expression. In contrast, NMDAR activation induced KCC2 internalization dependent on extracellular Ca^2+^ influx. In turn, calpain-mediated KCC2 degradation, but not internalization, required Ca^2+^ influx through voltage-gated Ca^2+^ channels. While TrkB-activation potentiated the effect of NMDAR on KCC2, the reverse was not true. Yet, strong NMDAR activation was sufficient to cause TrkB-independent KCC2 downregulation. Finally, prolonged, but not short-term inhibition of KCC2 activity caused NMDAR-dependent KCC2 downregulation. These findings reveal, for the first time, that a co-transporter function can be regulated through other means than membrane expression: through a continuum of interwoven synergistic processes, from function to internalization to degradation, scaling with time and stimulus strength.

## INTRODUCTION

Ionotropic *γ-aminobutyric acid type A* (GABA_A_) and glycine (Gly) receptor-mediated inhibition in the adult central nervous system (CNS) critically relies on transmembrane Cl^-^ flux^1^. A key molecule to define inhibitory synaptic strength is the neuron-specific K^+^-Cl^-^ cotransporter isoform 2 (KCC2) which normally extrudes Cl^-^ from the neurons to maintain a low intracellular concentration and compensate for Cl^-^ accumulation through GABA_A_ and Gly receptor/channels^2^. In conditions of on-going synaptic input, as is the case *in vivo*, which imposes a continuous Cl^-^load, a decrease in KCC2 function impairs the ability of the cell to maintain low intracellular Cl^-^. This leads to a weakening of GABA_A_ and Gly receptor-mediated inhibition, a deterioration of the robustness of inhibitory synaptic function and degradation of information transfer^3,4^. Consistent with this, KCC2 hypofunction has been proposed as a key substrate of several neurological and psychiatric disorders including epilepsy, motor spasticity, schizophrenia, chronic pain, neurodegenerative disorders, and drug-dependence^5–11^.

A major molecular pathway known to modulate KCC2 is the activation of tyrosine-receptor kinase B (TrkB) by its natural ligand brain-derived neurotrophic factor (BDNF). In mature neurons, BDNF/TrkB-signaling downregulates KCC2^12–14^. This appears as a substrate of several CNS disorders^5,12,14,15^. BDNF can be released from different sources, indicating multiple intercellular regulatory pathways. BDNF-TrkB signaling also appears to promote synaptic plasticity^16–22^, raising the question of whether this results from KCC2 downregulation. Similarly to BDNF/TrkB, N-methyl-D-aspartate (NMDA) receptors (NMDARs)/Ca^2+^ signaling pathway plays an important role in the modulation of Cl^-^ homeostasis as it has been shown to reduce KCC2 membrane clustering, plasmalemmal and cytoplasmic expression and to induce calpain-mediated KCC2 proteolysis in the CNS^23–26^. While these signaling pathways appear to affect KCC2 activity *via* transcriptional regulation^12,14^ or protein degradative mechanisms^5,12,14,23,24,27^, the question remains whether they affect function independent of expression.

To address this, and to explore how these two pathways interact, we stimulated TrkB and NMDAR individually and sequentially in rat cultured hippocampal neurons and in adult tissue. We found that TrkB signaling alone reduces both KCC2 function and protein in a ryanodine channel-dependent manner while Ca^2+^ influx through NMDA receptors induces KCC2 internalization. In turn, calpain-mediated degradation results from activation of voltage gated Ca^2+^ channel (VGCC). We also found that KCC2 modulation mediated by NMDAR signaling is potentiated by TrkB stimulation, but that the reverse was not true. Thus, these signaling mechanisms synergize, through mobilization of Ca^2+^ from different sources to regulate KCC2 function, internalization, and degradation.

## RESULTS

### Activation of TrkB signaling affects KCC2 function before expression

We first tested the effect of bath application of BDNF in the dorsal horn of adult spinal cord slices (SDH) on KCC2 function and expression. To assess transport, we measured transporter activity in reverse mode^28^ by measuring K^+^-driven uptake of Cl^−^ (Fig. 1a- c). We performed Cl^−^imaging and stepped extracellular K^+^ from 2.5 mM to 15 mM in control or BDNF-treated slices^6,7^. After 4h of BDNF treatment, Cl^-^ transport rate was significantly decreased compared with control (Fig. 1d; one-way ANOVA, *F* = 16.48, *p* < 0.0001, with post-hoc Tukey’s multiple comparisons, *p* = 0.0002). Co-treatment with the protease inhibitor cocktail Complete® did not prevent the decrease in transport rate (post-hoc Tukey’s multiple comparisons, *p* < 0.0001; Fig. 1d), indicating that change in Cl^-^ transport was independent of protein degradation. We then analysed spinal tissue treated for the same duration using a biotinylation protocol to isolate the membrane protein fraction (Supplemental Fig. 1). We found no significant change in surface (Wilcoxon matched-pairs, *W* = 51, *p* = 0.39), nor total KCC2 (*W* = 30, *p* = 0.60, Fig. 1e), indicating that BDNF signaling affected KCC2 function independent of protein trafficking or degradation.

**Figure 1:**
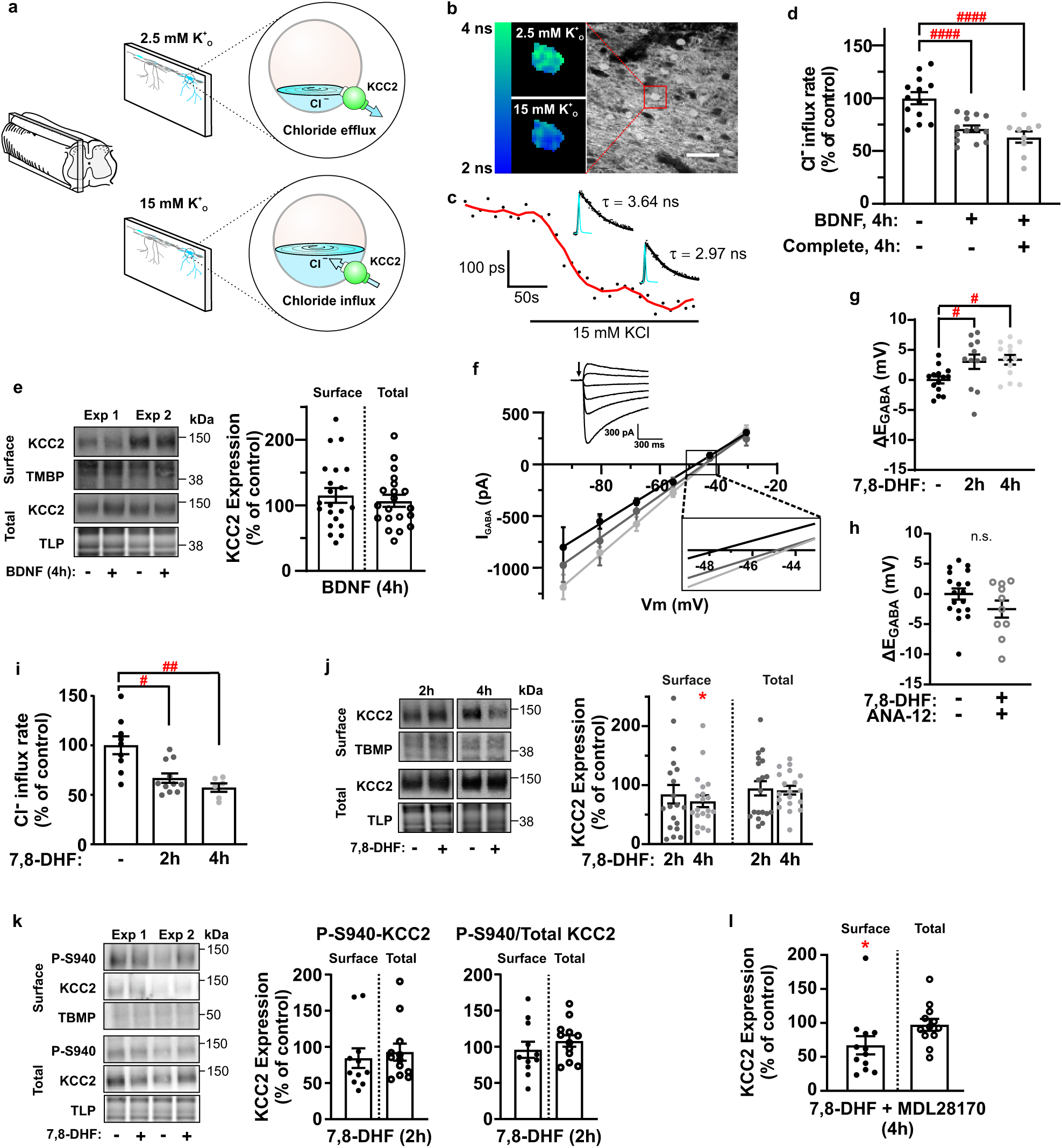
BDNF and 7,8-DHF downregulate KCC2 function without affecting expression. (**a**) Schematic representation of the protocol used to measure KCC2 activity in slices. (**b**) Color map (left) of the sum of 5 FLIM images from a control lamina II neuron (right) loaded with MQAE, before and after KCl. Scale bar 50 µm. (**c**) Trace of lifetime decay change of the neuron in (b) under KCl exposure. Inset: decay average before and after KCl. (**d**) Cl^-^ transport rate in SDH slices treated for 4h with BDNF (n = 9) or BDNF supplemented with the protein inhibitor cocktail Complete® (n = 14) as percentage of control mean. (**e**) Left: membrane and total fraction immunoblots of SDH slices treated or not with BDNF. Right: quantification of the surface and total KCC2 protein expression after BDNF compared with control (n = 21). (**f**) I-V relationships of representative control (black) and10 μM 7,8-DHF treated neurons for 2h (dark grey) and 4h (light grey). Inset: Representative trace of a lamina II neuron responding to puffs of 500 μM muscimol (arrow) at different holding voltages. (**g**) Pooled E_GABA_ of lamina II neurons treated with vehicle (●) or 10 μM 7,8-DHF for 2h (●) or 4h (●) (n = 14, 12, 13 neurons, respectively). Data are presented as the difference from control (DMSO) mean value. (**h**) Pooled E_GABA_ of lamina II neurons treated with vehicle (●) or 10 μM 7,8-DHF + 1 μM ANA-12 for >2h (○) (n = 18, 10 neurons, respectively). Data are presented as the difference from control (DMSO) mean value. (**i**) Cl^-^ transport rate in SDH control slices or treated with 7,8-DHF for 2h or 4h (n = 9, 11, 6 slices, respectively) as percentage on control mean. (**j**) Left: membrane and total fraction immunoblots of SDH slices treated or not with 7,8-DHF for 2h or 4h. Right: quantification of surface and total KCC2 protein expression after 7,8-DHF compared with control (2h: n = 19; 4h: n = 20). (**k**) Left: immunoblots from membrane and total fractions of P-S940 KCC2 from SDH slices treated or not with 10 μM 7,8-DHF for 2h or 4h. Middle: quantification of surface and total P-S940 KCC2 protein expression after 7,8-DHF compared with control (2h: n = 19; 4h: n = 20). Right: P-S940 to total KCC2 (within each fraction) ratio. (**l**) Quantification of the surface and total KCC2 protein expression after 7,8-DHF and the calpain inhibitor, MDL28170 (50 μM), compared with control (n = 12). TBMP = total biotinylated membrane protein; TLP = total lysate protein. Bars in the graphs represent the mean ± s.e.m. * and # = *p* < 0.05, ## = *p* < 0.01, #### = *p* < 0.0001.

Because BDNF does not only target TrkB but also p75^NTR^ receptors^29^, we repeated these experiments with the TrkB agonist 7,8-Dihydroxyflavone (7,8-DHF) ^30^. Using whole-cell patch-clamp recordings from spinal lamina II neurons with a high Cl^-^ intrapipette solution to impose a Cl^-^ load we found that 2h treatment of adult spinal cord slices with 7,8-DHF caused a significant depolarization of the reversal potential for GABA_A_ currents (*E*_GABA_; control: *E*_GABA_ = -48.08 ± 0.58 mV; 2h: *E*_GABA_ = -45.04 ± 1.2 mV; one-way ANOVA, *F* = 4.75, *p* = 0.015, with post-hoc Tukey’s multiple comparisons, *p* = 0.047) as reported for BDNF^15^, indicating that the Cl^-^ extrusion capacity was compromised by TrkB receptor activation (Fig. 1 f-g). We found no further depolarization of *E*_GABA_ with 4h of treatment (4h: *E*_GABA_ = -44.72 ± 0.81 mV; post-hoc Tukey’s multiple comparisons, control *vs.* 4h *p* = 0.022, 2h *vs.* 4h *p* = 0.96) indicating that a maximal functional effect was reached at 2h (Fig. 1g). Complementary Cl^-^ imaging led to similar findings (Kruskal-Wallis, *H* = 10.98, *p* = 0.0041; Fig. 1j). Moreover, incubation with the TrkB antagonist, ANA-12^31^, prevented the E_GABA_ depolarization induced by 7,8-DHF (unpaired *t*-test with Welch’s correction, *t* = 1.48, *p* = 0.16; Fig. 1 h), thus confirming the specific role of TrkB signaling in this mechanism.

In contrast to function, no difference in either surface or total KCC2 level were observed after 2h 7,8-DHF incubation (Wilcoxon matched-pairs: surface, *W* = -64, *p* = 0.21 and total, *W* = -40, *p* = 0.44; Fig. 1j), confirming TrkB-mediated modulation of function independent of trafficking. A decrease in KCC2 phosphorylation at serine 940 (P S940 KCC2) has been reported to cause KCC2 internalization^24,32^. Therefore, we tested whether the surface decrease that we observed at 4h was preceded by changes in P S940 KCC2. We found no change in surface or total P S940 KCC2 at 2h of 7,8 DHF incubation (Wilcoxon matched-pairs: surface, W = -24, p = 0.32 and total, W = - 30, p = 0.27; Fig. 1 k), nor in the ratio P S940 to total surface/total KCC2 (surface, W = -10, p = 0.7 and total, W = 18, p = 0.52; Fig. 1 k). This result indicates that the changes on KCC2 function observed at 2h were not due to onset of a signalling leading to internalization, but rather KCC2 functional modulation through other mechanisms.

In contrast to the 2h time point, after 4h, 7,8-DHF caused significant decrease in cell surface expression of KCC2 (Wilcoxon matched-pairs, *W* = -132, *p* = 0.012), without affecting the total expression (*W* = -52, *p* = 0.31; Fig. 1j). Blockade of calpain with MDL28170 did not prevent the decrease in surface expression of KCC2 at 4h (Wilcoxon matched-pairs, *W* = -54, *p* = 0.034; Fig. 1l), suggesting that it was due to internalization, not degradation, consistent with our previous results using the protease inhibitor cocktail (Fig. 1d).

### NMDARs activation decreases KCC2 expression

Previous studies have reported KCC2 downregulation through NMDAR stimulation^23–26,33^. Here we tested the effect of concentration and time of NMDA treatment on KCC2 expression. We found a dose- and duration-dependent downregulation of surface and total KCC2 (Fig. 2a, b). However, internalization appeared to precede total KCC2 downregulation (Fig. 2b). Based on the dose-response curve (Fig. 2a), 60 µM NMDA was the concentration used as a mid-range dose for the following experiments.

**Figure 2:**
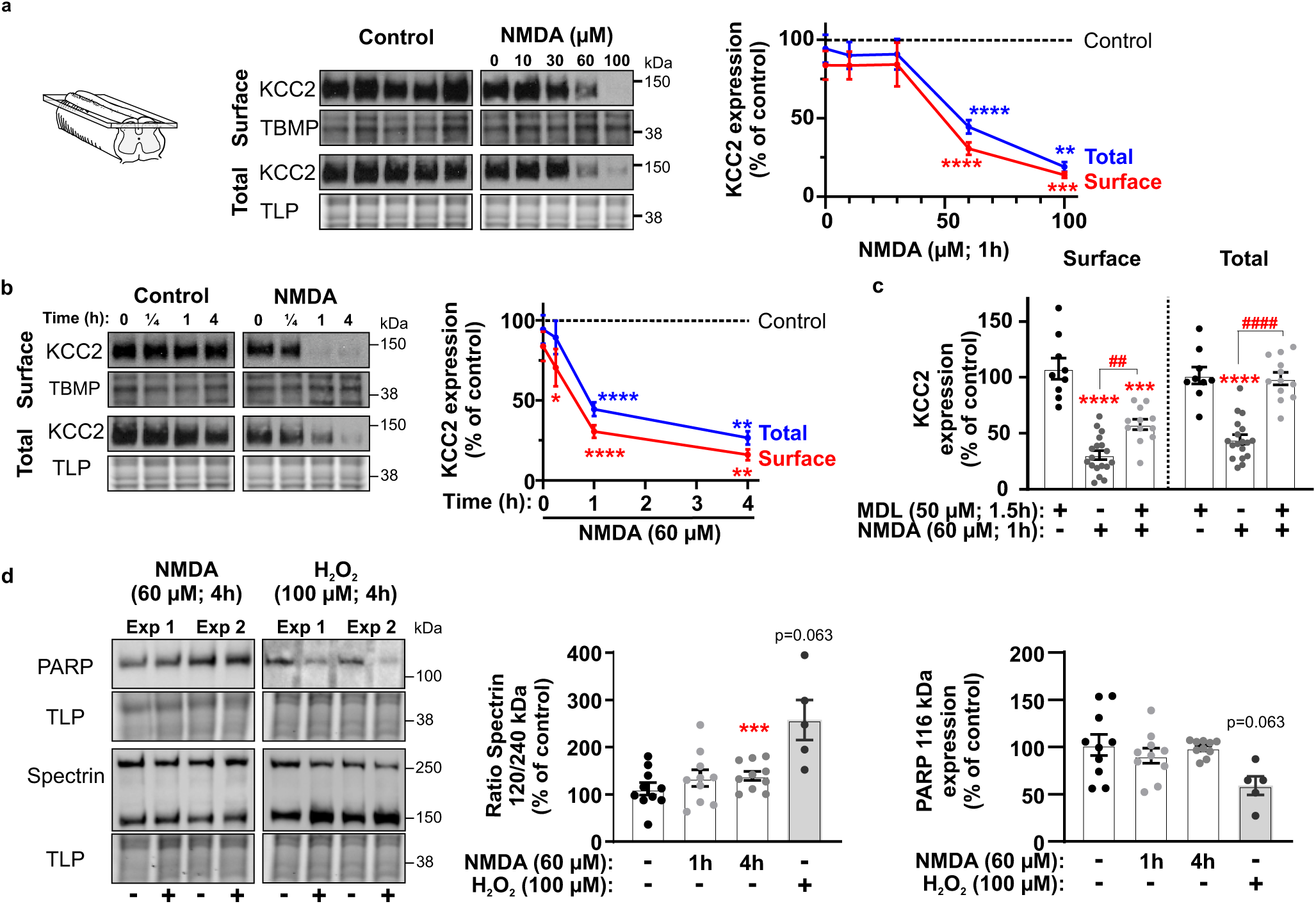
NMDAR stimulation sequentially induces internalization and degradation of KCC2. **(a**) Left: membrane and total fraction immunoblots of SDH slices (inset) treated with increasing concentrations of NMDA. Right: Dose-response curve of KCC2 surface and total protein expression to varying concentrations of NMDA for 1h compared with control (n = 8-19). (**b**) Left: membrane and total fraction immunoblots of SDH slices treated with NMDA for different time. Right: Time-response curve of KCC2 surface and total protein expression to NMDA compared with control (n = 10-19) (**c**) Quantification of the surface and total KCC2 protein expression after treatment with NMDA (1h) and/or the calpain inhibitor, MDL28170 (MDL, 1.5h), compared with control (n = 9-19). (**d**) Left : Spectrin and PARP immunoblots of SDH slices treated with NMDA or H_2_O_2_. Middle : Quantification of the cleaved vs. full length Spectrin protein levels. Left : Quantification of the full length PARP protein levels. TBMP = total biotinylated membrane protein; TLP = total lysate protein. Bars in the graphs represent the mean ± S.E.M. * = *p* < 0.05, ** and ## = *p* < 0.01, *** = *p* < 0.001, **** and #### = *p* < 0.0001.

Consistent with previous reports^23,26^, we found that co-treatment with the calpain inhibitor MDL28170 completely prevents the proteolysis of the total KCC2 protein levels (one-way ANOVA, *F* = 38.86, *p* < 0.0001, with post-hoc Tukey’s multiple comparisons, *p* < 0.0001; Fig. 2c). Remarkably, even though MDL28170 completely prevented total KCC2 downregulation, significant surface downregulation remained (Wilcoxon matched-pairs, *W* = -78, *p* = 0.0005), indicating distinct mechanisms for membrane *vs*. intracellular regulation of KCC2. This suggests that NMDA-dependent calpain activation is responsible for the proteolysis of the internalized KCC2 but not for KCC2 internalization itself.

NMDAR activation is a well-known substrate of neurotoxicity. To validate that our results are not mediated by neurotoxic cascades, we looked at spectrin, a well-known substrate for calpain proteolysis^34,35^. Spectrin cleaved vs. full length fraction (120/240 kDa) was significantly increased after 4h of NMDA treatment (Wilcoxon matched-pairs, *W* = -55, *p* = 0.002) but not after 1h (*W* = -33, *p* = 0.06; Fig. 2d). This supports that calpain is not mediating the membrane KCC2 decrease observed after 15 min of NMDA, but rather activated upon longer exposure. Moreover, this result also indicates that, at 1h, there is no significant cell death happening, yet some toxicity might be occurring after 4h. To assess whether the observations at 4h were caused by calpain activation or NMDA-triggered cell death, we looked at the level of poly (ADP-ribose) polymerase-1 (PARP), a DNA repair protein that is cleaved by caspases during apoptosis^36,37^. No significant change was observed in the non-cleaved 116 KDa PARP fraction under the same conditions (Wilcoxon matched-pairs: 1h, *W* = 27, *p* = 0.2; 4h, *W* = -4, *p* = 0.9; Fig. 2d), indicating that even if excitotoxicity is present after 4h, the changes were not triggered by cell-death mechanisms yet. In contrast, H_2_O_2_ pro-apoptotic conditions^38^ increased both spectrin cleavage and PARP degradation (Wilcoxon matched-pairs: Spectrin, *W* = -15, *p* = 0.03; PARP, *W* = 15, *p* = 0.03, Fig. 2d).

### TrkB signaling potentiates NMDAR-induced KCC2 protein downregulation

Since BDNF is known to potentiate NMDARs^39–42^, we hypothesised that pre-treating spinal slices with 7,8-DHF could potentiate the effect of NMDA on KCC2 downregulation. For total KCC2, while 15 min NMDA (Wilcoxon matched-pairs, *W* = -47, *p* = 0.15) and 4h 7,8-DHF (*W* = -52, *p* = 0.31) treatments had no effect on their own, the combined sequence of 7,8-DHF followed by NMDA (*W* = -162, *p* = 0.0061), but not the reverse sequence (*W* = -16, *p* = 0.57), caused significant downregulation (Fig. 3a), indicating a directional modulation of NMDA-mediated effects by TrkB signaling.

**Figure 3:**
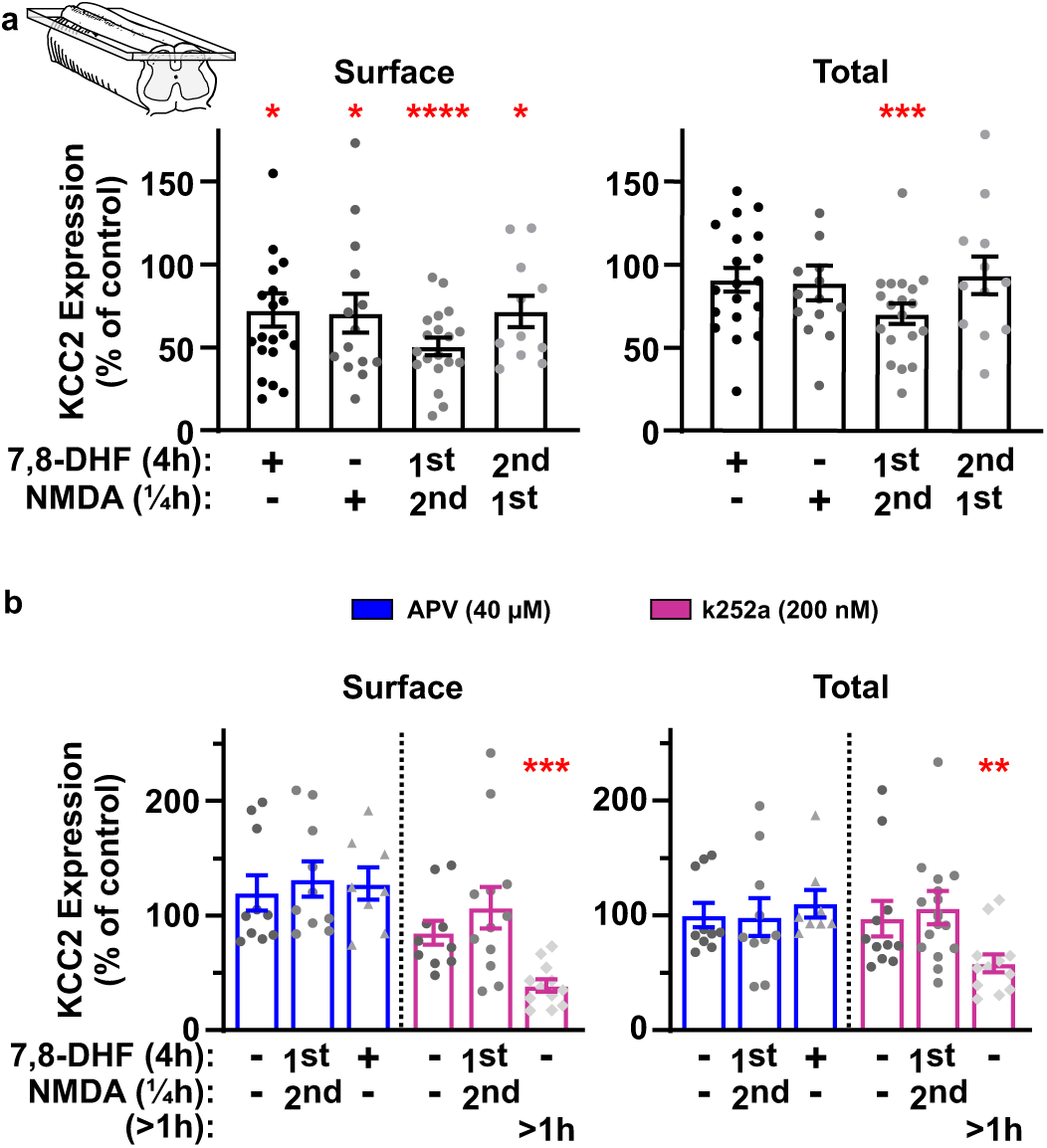
TrkB-signaling potentiates NMDAR-mediated downregulation of KCC2. **(a**) Quantification of the surface (left) and total (right) KCC2 protein expression on SDH slices (inset) after a subsequent incubation with 7,8-DHF and NMDA, and vice versa, or the two treatments alone, compared with control (n = 11-20). (**b**) Quantification of the surface (left) and total (right) KCC2 protein expression after a subsequent incubation with 7,8-DHF and/or NMDA in the presence of the NMDAR blocker, APV, or the TrkB inhibitor, K252a, compared with control (n = 8-12). Bars in the graphs represent the mean ± s.e.m. ** = *p* < 0.01, * = *p* < 0.05, *** = *p* < 0.001. One data point per panel is out of the graphs axis in (a) for graphical reasons but was included for the analysis.

The NMDAR antagonist (2*R*)-amino-5-phosphonovaleric acid (APV) prevented the decrease in membrane and total KCC2 induced by 7,8-DHF-potentiated NMDARs (Wilcoxon matched-pairs: surface, *W* = 22, *p* = 0.15, and total, *W* = -5, *p* = 0.85) and even the decrease in surface KCC2 induced by 7,8-DHF treatment alone (*W* = 29, *p* = 0.16; Fig. 3b). These results support that NMDARs are downstream effectors of TrkB-signaling for the internalization and degradation of KCC2. A slight increase in KCC2 surface expression was observed in all conditions where NMDA signaling was blocked by APV (Fig. 3b, left panel), suggesting that NMDARs may also be involved in the basal regulation of KCC2 membrane expression. The blockade of 7,8-DHF/TrkB signaling with the TrkB inhibitor K252a prevented the downregulation of KCC2 induced by 7,8-DHF-potentiated NMDARs (Wilcoxon matched-pairs: surface, *W* = -4, *p* = 0.91 and total, *W* = 4, *p* = 0.91), but K252a did not affect the KCC2 surface and total expression with longer NMDA incubation (surface, *W* = -78, *p* = 0.0005, and total, *W* = -72, *p* = 0.0024; Fig. 3b). These data show that 7,8-DHF/TrkB signaling potentiates NMDAR-signaling and is dependent of NMDAR pathway to modulate KCC2 protein levels, whereas NMDAR pathway *per se* can reduce KCC2 expression.

### Both TrkB and NMDA-signaling require intracellular Ca^2+^ rise for KCC2 downregulation

TrkB and NMDAR signaling pathways are known to mediate a rise in intracellular Ca^2+^ ([Ca^2+^]_i_). TrkB signaling can induce Ca^2+^ release from the endoplasmic reticulum through inositol 1,4,5-trisphosphate signaling^43^ and NMDAR activation mediates Ca^2+^ entry^44^. We tested the source and the role of Ca^2+^ in 7,8-DHF- and NMDAR-mediated KCC2 downregulation in the SDH. Co-application of 7,8-DHF and dantrolene, a blocker of intracellular Ca^2+^ release, prevented the surface KCC2 decrease mediated by TrkB signaling (Wilcoxon matched-pairs, *W* = 20, *p* = 0.41) while extracellular Ca^2+^ depletion did not prevent it (*W* = -57, *p* = 0.048; Fig. 4a, left panel). On the other hand, NMDA treatment had no significant effect on either surface or total levels of KCC2 when extracellular Ca^2+^ was depleted (surface, *W* = -29, *p* = 0.16, and total, *W* = -21, *p* = 0.32) but 20μM dantrolene did not prevent the NMDAR-mediated downregulation (surface, *W* = -28, *p* = 0.016, and total, *W* = -21, *p* = 0.016; Fig. 4a). These results highlight how Ca^2+^ signaling can affect KCC2 modulation differently and suggest that [Ca^2+^]_i_ levels could be the main trigger for the different mechanisms involved in the internalization and degradation of the cotransporter.

**Figure 4:**
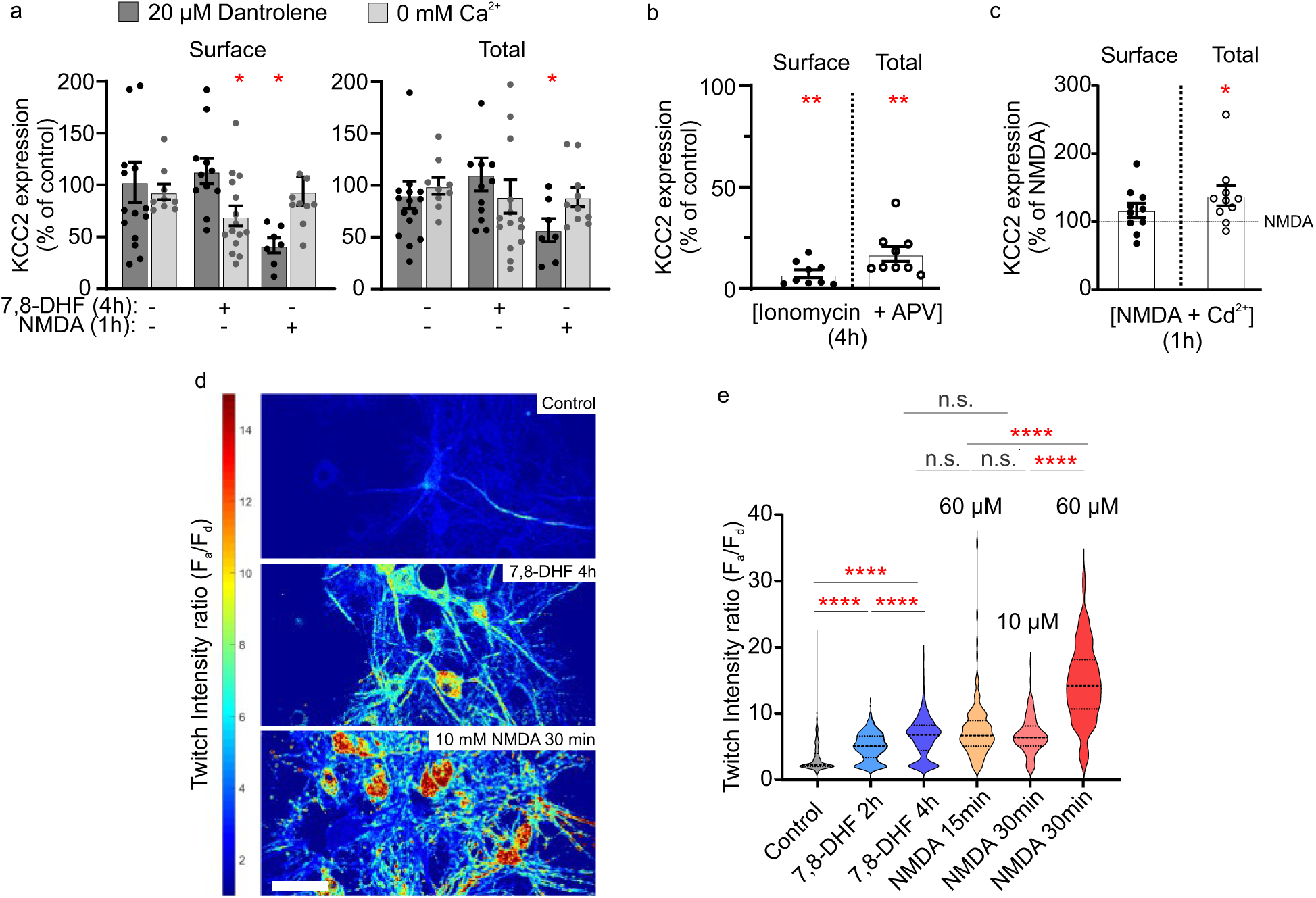
Differential KCC2 modulation relies on different Ca^2+^ sources. **(a**, **b, c**) Quantification of the surface (left) and total (right) KCC2 protein expression of SDH slices: (**a**) after incubation with 7,8-DHF or NMDA in the presence of the intracellular Ca^2+^ release blocker dantrolene (dark grey), or under extracellular Ca^2+^ depletion (light grey) compared with controls (n = 8-18); (**b**) After induction of Ca^2+^ influx with ionomycin in the presence of APV (n = 9), and (**c**) after incubation with NMDA supplemented or not with 10 μM cadmium (Cd^2+^; n = 10, 10). (B) Twitch-2b intensity ratio heatmap of hippocampal neuronal cultures expressing Twitch-2b in control conditions (top), after 4h with 10 μM 7,8-DHF (middle) and after 30 min with 10 μM NMDA (bottom). (**e**) Violin plot of average in Twitch-2b intensity ratio (ET540/40 channel over ET480/40 channel) from individual neurons in control conditions (n = 502), after 2h (n = 361) or 4h (n = 579) 7,8-DHF, NMDA 60 μM for 15 min (n = 161), 30 min (n = 93) and 10 μM for 30 min (n = 143). (**f**) Cumulative frequency distribution of the data in (e). Bars in the graphs represent the mean ± S.E.M. * = *p* < 0.05, **= *p* < 0.01, ### = *p* < 0.001, ****= *p* < 0.0001.

Application of the Ca^2+^ ionophore ionomycin in hippocampal slices reduces total KCC2 expression in a calpain-mediated mechanism^23^. We found that both surface and total KCC2 expression levels were strongly decreased in SDH slices treated with 50 μM ionomycin and 40 μM APV (Wilcoxon matched-pairs: surface, *W* = -45, *p* = 0.0039, and total, *W* = -45, *p* = 0.0039; Fig. 4b). Thus, a rise in [Ca^2+^]_i_ *per se* constitutes the central point in the cascade of events regulating KCC2 in adult tissue.

Since NMDAR activation unchains both internalization and degradation of KCC2, we tested whether NMDAR are the only path for Ca^2+^ entry involved in these mechanisms. To this end, we performed an NMDA treatment for 1h in the presence of 10 μM Cd^2+^, a voltage-gated Ca^2+^ channel (VGCC) blocker^40,45–48^ . We found that Cd^2+^ significantly reduced the degradation of total KCC2 (Wilcoxon matched-pairs: total, *W* = 49, *p* = 0.0098), but did not affect the membrane KCC2 internalization induced by NMDAR (*W* = 25, *p =* 0.23; Fig. 4c). This result indicates that NMDAR activation mediates a Ca^2+^ entry that is sufficient to induce KCC2 endocytosis but not degradation. VGCCs activation, secondary to NMDAR signaling, is necessary to trigger degradation.

Since different Ca^2+^ pools appear to be at the root of the differential regulation of KCC2, we explored whether this was also associated with different [Ca^2+^]_i_. To be able to compare steady-state Ca^2+^ levels across cells, we transfected hippocampal cultures with a virus expressing the genetically encoded ratiometric Ca^2+^ sensor, Twitch-2b^49^, and imaged the neurons at 19-23 DIV. Incubation with 10 µM 7,8-DHF induced a gradual increase in Twitch-2b intensity ratio, indicating a progressive rise in [Ca^2+^]_i_ at 2h and 4h (Kruskal-Wallis, *H = 605.2*, *p < 0.0001,* Fig. 4 d-e). NMDA treatment also increased [Ca^2+^]_I_, reaching, within 15 min, similar levels to that at 4h of 7,8-DHF treatment (post-hoc Dunn’s multiple comparison test, 4h 7,8-DHF vs. 15 min NMDA 60μM, *p* = 0.61). The NMDA-dependent increase in Ca^2+^ was dose and time dependent (post-hoc Dunn’s multiple comparison test: control vs. 15 min NMDA 60 μM, *p* < 0.0001; control vs. 30 min NMDA 60 μM, *p* < 0.0001; control vs. 30 min NMDA 10 μM, *p* < 0.0001; Fig. 4 d-e). Therefore, different Ca^2+^ levels appear to account for the different regulation of KCC2 at the functional, trafficking and degradation levels, respectively.

### Prolonged TrkB signaling engages NMDAR for KCC2 degradation

The membrane decrease observed after 4h of treatment with 7,8-DHF (Fig. 1k) raised the question of the role of 7,8-DHF/TrkB signaling pathway in the control of KCC2 protein levels. We explored the effects of longer exposures to 7,8-DHF on the modulation of KCC2. As SDH slices cannot survive for long periods, we used primary hippocampal cultures for these experiments. We first corroborated that 4h of 7,8-DHF had the same effect in cultures than in SDH slices (Fig. 5a, b). We also found a decrease in the surface levels of KCC2 after 4h (Wilcoxon matched-pairs, *W* = -128, *p* = 0.0002), while no change was found in the total protein level (*W* = -62, *p* = 0.12; Fig. 5b). However, longer incubation with 7,8-DHF produced a progressive KCC2 downregulation at both, membrane (24h, *W* = -101, *p* = 0.0004, and 7 days, *W* = -78, *p* = 0.0005) and cytosolic levels (24h, *W* = -73, *p* = 0.02, and 7 days, *W* = -68, *p* = 0.0049; Fig. 5b). Interestingly, the KCC2 degradation after either 24h or 7 days of 7,8-DHF treatment was prevented with APV during the same period (unpaired t-test: 24h, *t* = 0.98, *p* = 0.34, and 7 days, *t* = 0.91, *p* = 0.37; Fig. 5c). These results confirmed that degradation of KCC2 by prolonged TrkB signaling requires NMDAR activation through endogenous glutamate.

**Figure 5:**
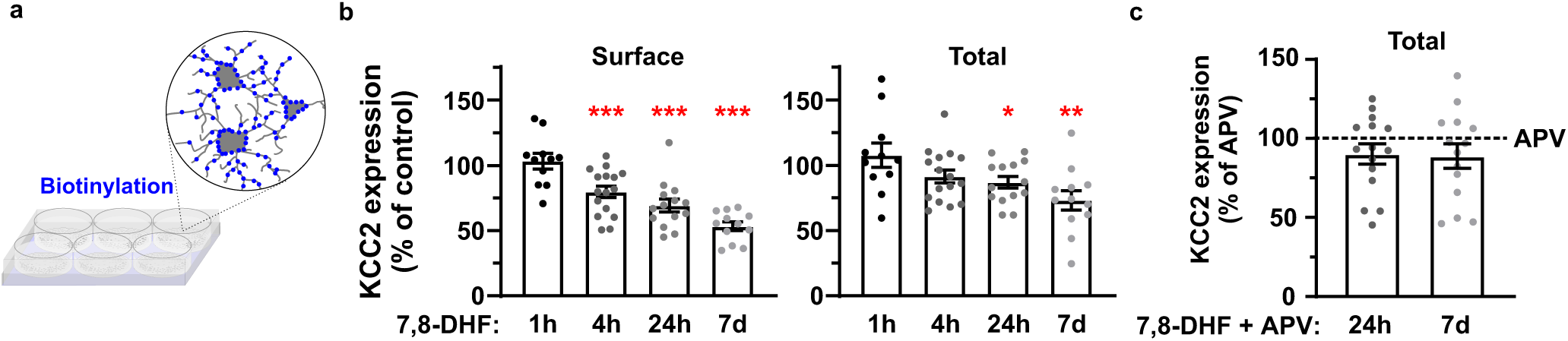
Long term TrkB-signaling induces a progressive KCC2 internalization and NMDAR-dependent degradation. (**a**) Schematic representation of the cell surface biotinylation on rat primary hippocampal cultures used for long term exposure KCC2 measurements. (**b**) Quantification of KCC2 surface (left) and total (right) protein levels after exposure to 7,8-DHF for 1h, 4h, 24h and 7 days compared with control (n = 11-16). (**c**) Quantification of total KCC2 protein levels after incubation with 7,8-DHF and APV for 24h or 7 days as percentage of APV treated cultures mean (n = 15, Unpaired t-test). Data are present in the graph as a percentage of the mean value of the only APV samples. (c) Quantification of total KCC2 protein levels after incubation with 7,8-DHF and APV for 24h or 7 days as percentage of APV treated cultures mean (n = 15, Unpaired t-test). Data are present in the graph as a percentage of the mean value of the only APV samples. Bars in the graphs represent the mean ± s.e.m. *= *p* < 0.05, **= *p* < 0.01, ***= *p* < 0.001.

### Maintained KCC2 inhibition prompts NMDAR-dependent KCC2 downregulation

Our previous results strongly suggest that the downregulation of KCC2 is a consequence of a rise in Ca^2+^. Disinhibition is known to potentiate NMDARs in the SDH^39,40^ and may generate a rise in Ca^2+^ due to hyperactivity. We tested the effect of blocking KCC2 function with the selective inhibitor VU0240551 to induce a disinhibition. No effect of VU0240551 was found on KCC2 expression in SDH slices after 4h (Wilcoxon matched-pairs: surface, *W* = 19, *p* = 0.58, and total, *W* = -9, *p* = 0.81, Fig. 6a). With longer exposure (24h) to VU0240551 in cultures we found a decrease in both, surface (*W* = -136, *p* < 0.0001) and total (*W* = -136, *p* < 0.0001) levels of KCC2 that was prevented by APV (surface: Kruskal-Wallis, *H* = 30.09, *p* < 0.0001, with post-hoc Dunn’s multiple comparison test, *p* < 0.0001; total: *H* = 29.17, *p* < 0.0001, post-hoc *p* < 0.0001; Fig. 6b). No significant difference was found between APV and APV+VU0240551 treated cultures in either surface or total KCC2 levels (post-hoc Dunn’s multiple comparison: surface, *p* > 0.99; total: *p* > 0.99). 24h of APV treatment in hippocampal cultures generated a high increase in KCC2 at both, plasmalemmal and total KCC2 level (Wilcoxon matched-pairs: surface, *W* = 105, *p* = 0.0001, and total, *W* = 103, *p* = 0.0002; Fig. 6b), which was independent of an effect of 24h APV on overall membrane protein expression (Supplemental Fig. 2). Further experiments would be necessary to elucidate the role of NMDAR on basal KCC2 regulation. Nevertheless, these results show that disinhibition by blockade of KCC2 eventually causes KCC2 downregulation over the long term and that this is dependent on NMDARs.

**Figure 6:**
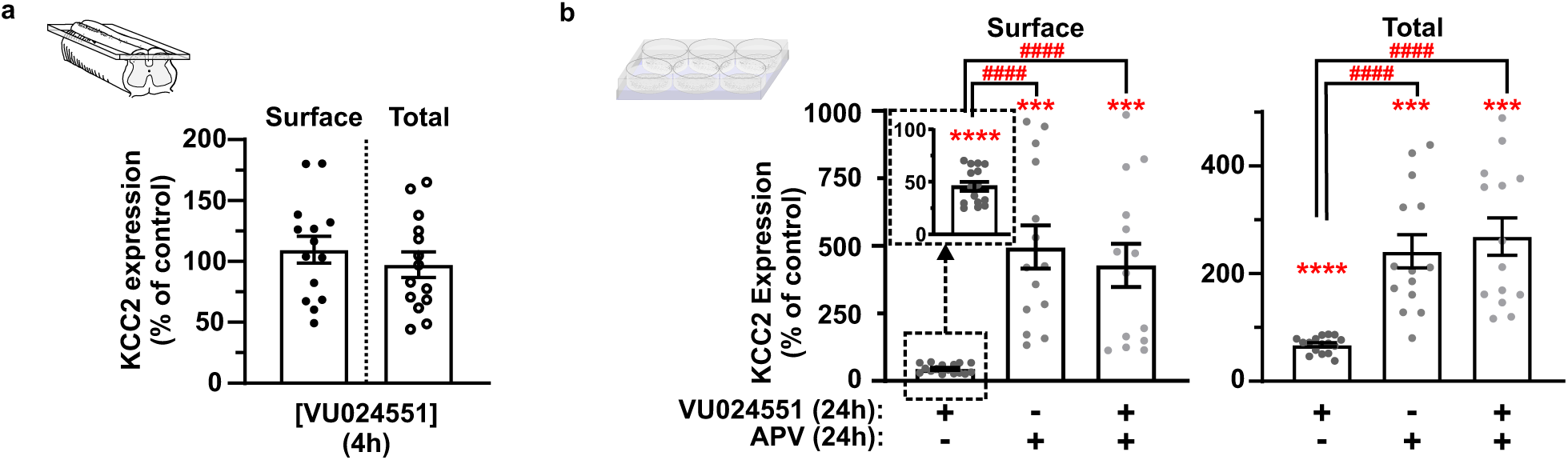
Long term disinhibition induces KCC2 downregulation. (**a**) Quantification of KCC2 surface and total protein levels in SDH slices (inset) after 4h incubation with the KCC2 antagonist VU024551 compared with control (n = 14). (**b**) Quantification of KCC2 surface and total protein levels in hippocampal cultures (inset) after 24h incubation with the KCC2 antagonist VU024551 and/or APV, compared with control (n = 14-16). Left insets in (a) and (b) represent the experimental system used for each panel. Bars in the graphs represent the mean ± s.e.m. *** = *p* < 0.001, ****and #### = *p* < 0.0001.

### Activation of TrkB signaling affects KCC2 function via ER-released Ca^2+^ but independently of NMDAR and surface protein levels

Our previous results suggest that 7,8-DHF effect may have a NMDAR-independent component acting specifically on KCC2 function. To test this hypothesis directly, we performed Rb^+^ influx assays in *Xenopus laevis* oocytes, a system that is regularly used to overexpress and study the ionic transport of cation-chloride cotransporters^7,50^. At stage V-VI, oocytes do not express TrkB receptor endogenously, so we co-injected TrkB and KCC2 cRNAs to activate TrkB-dependent intracellular event cascades (Fig 7a); a strategy previously used^51^. We found a significant effect of both cRNAs expressed and drug effects: coexpression of TrkB and KCC2 (TrkB + KCC2) generated higher Rb^+^ intake in the oocytes compared to the ones where only TrkB was overexpressed (TrkB), confirming significant expression of KCC2 (two-way ANOVA, *F*_cRNA_ = 24.89, *p* < 0.0001; Fig. 7b) and 7,8-DHF treatment decreased the Rb^+^ influx rate by a 40-45% in TrkB + KCC2 oocytes but not in TrkB oocytes (*F*_treatment_ = 24.89, *p* < 0.0001; *F*_interaction_ = 13.11, *p* = 0.0004; post-hoc Sidak’s multiple comparisons: control *vs*. 7,8-DHF in TrkB oocytes *p* < 0.0001, and in TrkB + KCC2 oocytes *p* < 0.0001; Fig. 7b). The 7,8-DHF dose-dependently decreased KCC2 transport activity in oocytes (IC_50_ of 47.95 µM; Fig. 7c). All together these data validate the oocyte heterologous expression system to study KCC2 functional regulation by TrkB-signaling.

**Figure 7:**
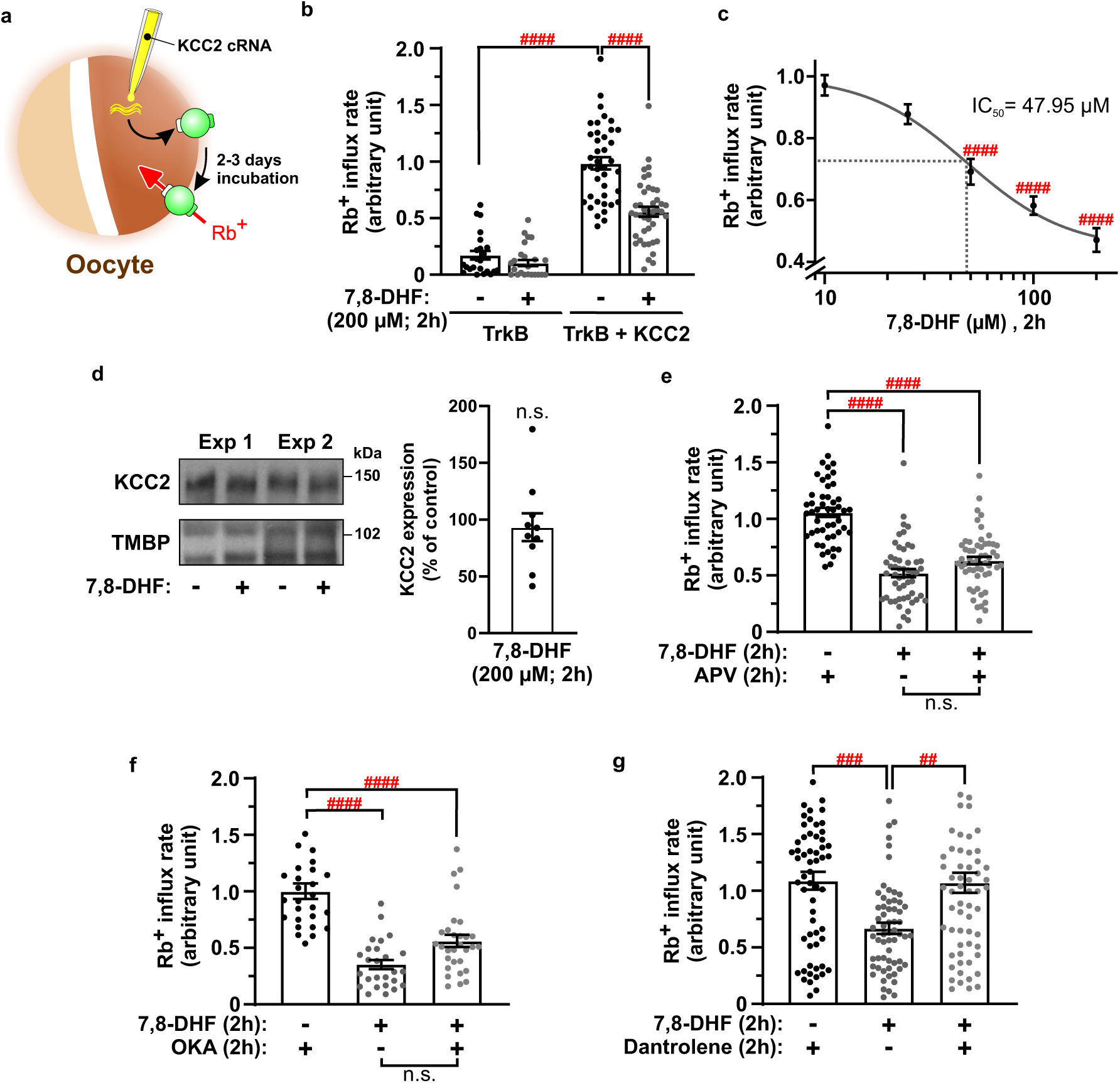
TrkB-signaling requires intracellular Ca^2+^ release but it is independent of NMDARs. **(a**) Schematic representation of the protocol used in the Rb^+^ assays with *Xenopus laevis* oocytes. (**b**) KCC2 transport activity measured through Rb^+^ influx rates after treatment with 7,8-DHF or vehicle of oocytes injected with only TrkB or TrkB + KCC2 cRNA. (**c**) Dose-response curve of the Rb^+^ influx rate of TrkB + KCC2 oocytes exposed for 2h to increasing concentrations of 7,8-DHF. (**d**) Left: Immunoblot of surface KCC2 expression of two batches of oocytes incubated with DMSO or 200 μM 7,8-DHF for 2h. Right: Quantification of surface KCC2 surface expression of oocytes incubated with 7,8-DHF relative to control (DMSO). (**e-g**) Rb^+^ influx rates of TrkB + KCC2 oocytes treated with 7,8-DHF and/or APV to block NMDARs signaling (**e**), okadaic acid (OKA) to block protein phosphatase 1-dependent internalisation (**f**), and dantrolene to block intracellular Ca^2+^ release (**g**). Bars in the graphs represent the mean ± s.e.m. ## = *p* < 0.01, ### = *p* < 0.001, #### = *p* < 0.0001, n.s. = not significant.

To ensure that the 7,8-DHF-induced KCC2 function deficit in oocytes is not due to a decrease in KCC2 cell surface expression, we measured the protein expression of membrane KCC2 on TrkB + KCC2 oocytes. The data indicate that the decline in Rb^+^ transport within the TrkB + KCC2 oocytes treated with 7,8-DHF is not associated with a decrease in membrane KCC2 (Wilcoxon matched-pairs: *W* = -19, *p* = 0.38; Fig. 7d). Additionally, APV did not prevent the effect of 7,8-DHF on KCC2 transport activity in oocytes (Kruskal-Wallis, *H* = 69.9, *p* < 0.0001, with post-hoc Dunn’s multiple comparisons: APV *vs*. APV + 7,8-DHF, *p* < 0.0001; Fig. 7e). Protein phosphatase 1 (PP1) is known to dephosphorylate KCC2 at S940, triggering the internalization^24^. However, treatment with a ∼10X saturating dose of okadaic acid (OKA)^50^, an antagonist of PP1, also did not prevent the decrease in the Rb^+^ intake mediated by 7,8-DHF (Kruskal-Wallis, *H* = 42.8, *p* < 0.0001, with post-hoc Dunn’s multiple comparisons: OKA *vs*. OKA + 7,8-DHF, *p* < 0.0001; Fig. 7f). These observations indicate that TrkB-signaling can affect KCC2 function independently of NMDAR activity, PP1-dependent dephosphorylation or trafficking.

We have previously shown the importance of Ca^2+^ in regulating KCC2 protein expression and trafficking (Fig. 4). To test for a role of Ca^2+^ in TrkB-mediated regulation of KCC2 function, we first blocked the release of Ca^2+^ from the ER with dantrolene in our oocyte transport assay. We found that it prevented the 7,8-DHF-induced KCC2 decrease in function (Kruskal-Wallis, *H* = 18.22, *p* = 0.0001, with post-hoc Dunn’s multiple comparisons: 7,8-DHF *vs*. dantrolene + 7,8-DHF, *p* = 0.0021; Fig. 7g). Thus, a release of Ca^2+^ from intracellular stores is necessary for TrkB-mediated regulation of KCC2 function. These results also reveal a role of Ca^2+^ not only in the control of KCC2 expression but also its transport activity.

## DISCUSSION

Regulation of KCC2 has emerged as a critical factor in the etiology of several neurological and psychiatric disorders^4–6,8,9,52–55^. Most studies to date have tended to equate changes in Cl^-^ transport to altered KCC2 expression and trafficking^12,14,24,26,32,53,56^. Yet, some studies report that phosphorylation of certain residues on KCC2 affect its function with no apparent effect on trafficking^57,58^, indicating that not all regulation necessarily occurs through trafficking only. Vale at al. reported changes in KCC2 function, independent of protein levels in the inferior colliculus after deafness^59^, but they base their quantification on total KCC2 measurements, precluding surface vs. total quantification. In our study, by quantifying changes in both compartments, we were able to show that TrkB receptor activation, by itself, in absence of NMDA receptor activation, affects function without affecting trafficking. Some studies report faster changes in KCC2 protein levels than the ones we observed^14,32^, but it was suggested that this was due to non optimal tissue conditions, resulting in heightened calpain activity^23^. By maintaining optimal conditions to preserve slice viability, in our study we were able to temporally separate functional from trafficking changes.

There have been several studies reporting downregulation of KCC2 as a consequence of TrkB-and/or NMDAR-signalling, as well as calpain-dependent degradation in rodents^15,23–26,60,61^ but also in human tissue^39^ . Our study now reveals that these mechanisms are only indirectly coupled. TrkB-mediated internalization occurs necessarily through NMDAR-signalling and S940 phosphorylation, secondary to TrkB activation. KCC2 degradation is also secondary to TrkB-and/or NMDAR-signalling as it requires VGCC activation. Our findings thus reconcile previous findings under a common hierarchical organization: from TrkB-mediated functional changes on a short time scale to NMDAR-mediated trafficking and, eventually, VGCC-mediated degradation (Fig. 8a, b). We also found that the hierarchical organization of mechanisms responsible for KCC2 regulation can be explained by different forms of Ca^2+^ mobilization: from TrkB-mediated release from intracellular stores to Ca^2+^ entry through NMDARs and further entry through VGCCs (Fig. 8a, b). This new framework forces a re-analysis of previous data through different lenses. For example, this can easily explain how conditions causing network hyperexcitability are associated with KCC2 internalization, through heightened activity-dependent NMDAR activation^14,23–25,53^. In turn, degradation will occur in any conditions leading to excitotoxic activity.

**Figure 8:**
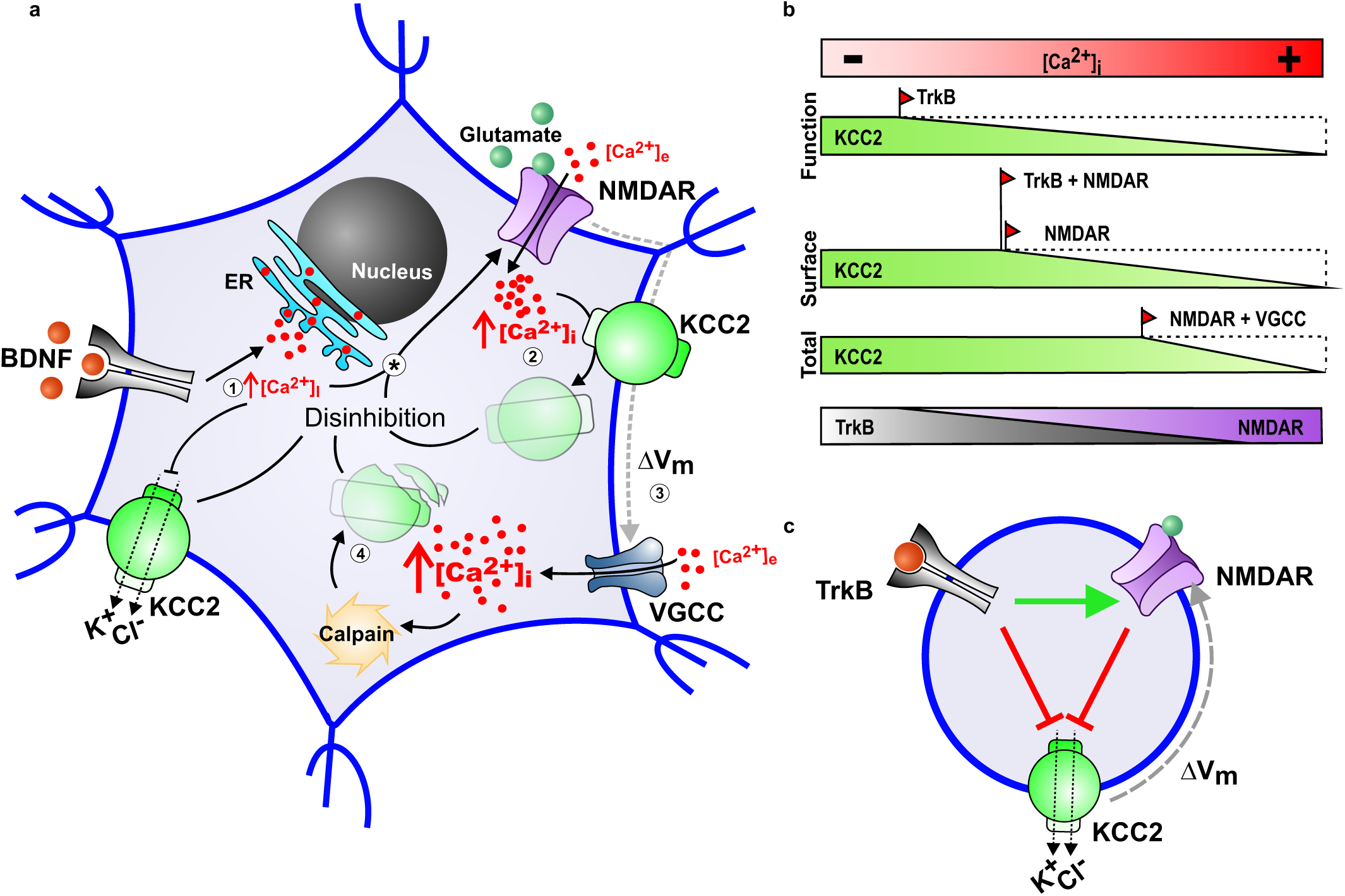
Schematic diagram of TrkB- and NMDAR-dependent post-translational regulation of KCC2. **(a**) 1, Increases in BDNF release in the SDH activate neuronal TrkB receptors which induce an intracellular store Ca^2+^ release. This Ca^2+^ release is responsible of a downregulation of KCC2 function; 2, Excitotoxic concentrations of glutamate acting on NMDARs induce a [Ca^2+^]_I_ rise that will mediate the endocytosis of KCC2; 3, NMDARs activation also mediate a membrane depolarization that will open VGCC and consequently a bigger [Ca^2+^]_i_ increase; 4, The Ca^2+^-sensitive protease calpain activates upon the higher [Ca^2+^]_i_ levels and degrade KCC2. All these mechanisms will result in a global KCC2 dysfunction in the SDH neurons, and general disinhibition. Both TrkB-signaling and the disinhibition will in turn potentiate NMDAR (*) ^39,40^. (**b**) The gradients in the diagram show how the different mechanisms modulating KCC2 (functional downregulation, endocytosis and degradation; *green gradients*) are triggered by [Ca^2+^]_i_ (*red gradient*) in SDH neurons. When [Ca^2+^]_i_ increases above the physiological levels, the activity of KCC2 is rapidly compromised. Progressive [Ca^2+^]_i_ rises first activate internalization pathways that result in less membrane KCC2 available. Finally, as [Ca^2+^]_i_ keeps increasing, the proteolytic mechanisms end up degrading the total levels of KCC2. TrkB (*grey gradient*) activation is sufficient to trigger this signaling loop since, besides the downregulation of KCC2 extrusion capacity, it will potentiate NMDARs (*purple gradient*). A higher NMDAR activity will then stimulate KCC2 internalization and later degradation. On the other hand, excitotoxic activation of NMDAR will prompt KCC2 endocytosis and degradation even in the absence of TrkB signaling due to the high [Ca^2+^]_i_ rises of this kind of activation. (**c**) Disinhibition potentiates NMDARs, connecting TrkB- and NMDAR- signaling pathways in a feedforward loop.

A key finding from our results is the functional regulation of KCC2 following TrKB signaling, independent of changes in trafficking. KCC2 membrane clusters are commonly associated with synapses^62–64^ and appear to be dynamically regulated by neuronal activity^25,65^. Excitatory activity increases KCC2 diffusion through NMDAR activation and Ca^2+^ influx^25^. Moreover, this increase in KCC2 diffusion with excitatory activity was observed when endocytosis was blocked, indicating that enhancement in KCC2 diffusion precedes that of KCC2 internalization^25^. In a recent study, Dedek and colleagues found a decrease in KCC2 synaptosomal levels after 70 min of BDNF incubation^39^, a result that may appear to contradict our findings. However, if KCC2 diffuses away from synaptic clusters, this may be detected as a decrease in KCC2 within synaptosomes but not necessarily within the whole membrane, as we measure, unless internalization is engaged.

Supporting this idea, Donneger et al. recently found that KCC2 activity could be enhanced using pharmacological approaches in a mechanism that requires reduced KCC2 membrane diffusion and increased clustering, but that is independent of membrane KCC2 levels^66^ . Taken together, these findings suggest that increased KCC2 membrane diffusion may be a mechanism responsible for TrkB-mediated functional downregulation, independent of internalization.

Recent studies show that TrkB-signaling, combined with disinhibition causes potentiation of NMDARs via the release of Ca^2+^ from intracellular stores^39,40^. Here we found that the TrkB-mediated release of Ca^2+^ from intracellular stores also affects KCC2 function. The combination of these two phenomena can set the stage for a positive feedback loop, compounding the collapse of inhibition through synergistic signalling cascades (Fig. 8c). On short time scales, however, both data from Hildebrand et al. and ours are in accord to say that blocking KCC2 alone is not sufficient to potentiate NMDAR^40^, nor to cause ensuing KCC2 downregulation (our results). Yet, we found that over the course of 24h, KCC2 blockade can eventually lead to a collapse in KCC2 in an NMDAR-dependent manner. We surmise that the heightened network activity, through prolonged disinhibition, eventually causes NMDAR potentiation^39,40^ (Fig. 8c). Over time, loss of KCC2 function may thus be sufficient to trigger a catastrophic collapse in inhibition^3,10^.

Our study features an apparently canonical time- and Ca^2+^-dependent sequence in KCC2 regulation, that occurs through an interwoven continuum of synergistic cellular processes, from function to trafficking to degradation, and which may delineate a fine balance between physiological ionic plasticity and pathophysiological disinhibition (Fig. 8). Our findings may pave the way to discovery of therapeutic targets that would spare a normal, physiological role for Cl^-^-mediated metaplasticity^4,67–69^.

## ACKNOWLEDGEMENTS

We thank Sylvain Côté for expert assistance with the illustrations and Dr. Paul Isenring for allowing use of his atomic absorption spectrometer 240Z AA (Agilent Technologies). We also thank Francine Nault and the Neuronal Cell Culture Platform of the Neurophotonics Centre, funded by the Canadian Neurophotonics Platform, for producing and maintaining neuronal cultures. The work was supported by CIHR grant FDN-159906 and a Simons Foundation grant to Y.D.K.. Y.D.K. was supported by a Canada Research Chair. A.G.G. is a Chercheur-Boursier (Fonds de recherche du Québec – Santé) and was supported by a Sentinel North Partnership Research Chair. M.J.B. was supported by a fellowship of the Savoy Foundation. M.C. was supported by a postdoctoral fellowship from the Fonds de Recherche du Québec – Santé.

## AUTHOR CONTRIBUTIONS

M.J.B., I.P-F., A.G.G. and Y.D.K. conceived and designed the experiments. M.J.B., I.P-F., A.B., N.C. and M.C. performed the experiments; M.J.B., I.P-F., A.B., M.C. and A.G.G. analyzed the data. I.P-F, M.J.B. and Y.D.K. wrote the manuscript.

## Competing interest

The authors declare no competing interest.

## METHODS

### Animals

All procedures were conducted in accordance with the guidelines of the Canadian Council on Animal Care and approved by the Université Laval Animal care committee. Male Sprague-Dawley CD rats (>90 day-old) were used for the experiments. The animals were housed in a room with an inverse 12h light/dark cycle.

### Primary neurons

Rat hippocampal neuronal cultures were prepared as previously described ^70^. Briefly, rat hippocampi were dissected out of post-natal day 0 (P0) rats and cells were dissociated both enzymatically using 12 U/ml papain (Worthington Biochemical Corporation) and mechanically by trituration with a Pasteur pipette. After dissociation, cells were centrifuged and plated on poly-d-lysine coated 3.5 cm plates (1x10^6^ cells/plate) or plated poly-d-lysine coasted 12 mm coverslip (Neuvitro) at 3x10^4^ cells/coverslip, and maintained in Neurobasal medium supplemented with B27 (50:1), 50 U/ml penicillin/ 50 µg/ml streptomycin mixtures, 0.5 mM L-glutamax and 2% FBS-HI. To reduce the number of glial cells, 5 μM of cytosine arabinofuranoside (ARA-C; Sigma) was added 5 days after plating and cells were kept in culture for 5 additional days in the presence of ARA-C. The day after and twice a week thereon, half of the growth medium was replaced with medium without ARA-C. Neurons were plated in 6-well plates (1M cells/well). Neurons were propagated at 37°C in a humidified atmosphere of 95% air and 5% CO_2_. Since KCC2 increases with maturation ^71^, we used cultures from 21 days *in vitro* to make sure KCC2 levels were already high in those neurons.

### Acute slice preparations

Acute spinal cord slices were prepared as previously described ^72^. Briefly, adult male rats were anesthetised with ketamine/xylazine and perfused with ice-cold slicing solution containing (in mM) 252 sucrose, 10 D-glucose, 2.5 KCl, 1.5 CaCl_2_, 6 MgCl_2_, 40 kynurenic acid (bubbled with 95% O_2_/ 5% CO_2_, pH 7.4). The animals were decapitated, and the spinal cord was extracted by hydraulic extrusion. Horizontal (250 µm thick; for protein assessment) and parasagittal (350-400 μm thick; for functional assessment) slices of the lumbar part of the spinal cord were cut in ice-cold slicing solution using a Leica VT1200S microtome. Slices were then incubated for a recovery period (15-60 min; 34°C) in artificial cerebrospinal fluid (ACSF) containing (in mM) 126 NaCl, 26 NaHCO_3_, 10 glucose, 2.5 KCl, 2 CaCl_2_, 2 MgCl_2_, 1.25 NaH_2_PO_4_ (bubbled with 95% O_2_/ 5% CO_2_, pH 7.4). For immunoblotting experiments, after recovery, horizontal slices were separated in two equal pieces along the midline and one of the sides was always used as control while the other was subjected to the specific treatments.

### Treatments

Rat hippocampal cultures and/or SDH slices were incubated for different durations in: 50 ng/ml BDNF (R&D Systems); 10 µM (SDH and cultures) or 10, 25, 50, 100 or 200 µM 7,8-DHF (oocytes; Sigma); 1 µM ANA-12 (Sigma); 10, 30, 60, and 100 µM NMDA (Tocris); 50 µM MDL28170 (Sigma); 40 µM APV (Tocris); 200 nM K252a (Sigma); 20 µM dantrolene (Sigma); 50 µM ionomycin (EMD Millipore); 200 µM cadmium (Sigma); 100 µM VU0240551 (Tocris); 1 µM okadaic acid (Sigma) and 1 pill of Complete® (Roche) in 40 mL of solution.

### Imaging of Cl− transport

Parasagittal lumbar SDH slices (400 μm thick) were kept before imaging for a maximum of ∼5.5h in bubbled ACSF at 34 °C. During this period slices were treated with DMSO, BDNF or 7,8-DHF for 2h or 4h. Slices were then loaded with the Cl^-^ indicator MQAE (N-6-methoxyquinolinium acetoethylester, Molecular Probes) as in previously reported ^6,7^. Briefly, slices were incubated in ACSF containing 5 mM MQAE for 30–40 min and transferred to a perfusion chamber where the extracellular MQAE was washed out for 10 min in the presence of 10 µM 6-cyano-7-nitroquinoxaline-2,3-dione (CNQX), 40 µM APV, 1µM strychnine and 1 µM tetrodotoxin (TTX) and 10 μM gabazine to minimize KCC2-independent Cl^−^ transport. Lamina II cells were visually identified by merging transmitted light and MQAE fluorescence. Lifetime imaging of MQAE was conducted as previously described ^3,6,7^. Two-photon excitation at 750 nm was used combined with a bandpass emission filter (475/30 nm) to block the laser during the acquisition. Instrument response function of the detection path was acquired using an 80-nm gold nanoparticle suspension to generate a second-harmonic signal. Lifetime images were acquired for 3s every 10s. After a control period of 50s, perfusion solution was switched to ACSF containing 15 mM KCl to reverse KCC2-mediated Cl^−^ transport (Fig. 1a, b, c). Lifetime in each cell was averaged over the whole soma and extracted for each time point using custom-made Matlab software. Lifetime changes for each slice were then expressed as the mean of changes occurring in each cell. Briefly, based on the work of Digman et al. ^73^, we converted the photon timing histograms of each acquired lifetime image to phasor plots. Then, for every time-point, regions of interest (ROIs) corresponding to cell bodies were manually selected and added to a new phasor. Lifetimes obtained for the cells were averaged for each slice at each time point to generate the lifetime time course. Chloride influx rates were then calculated between the initial and final lifetime plateaus.

### Electrophysiological recordings

Whole-cell patch clamp recordings were performed in 350 μm thick parasagittal slices of lumbar spinal cord from male rats. Slices were transferred to regular ACSF (previously described) after slicing and incubated at 34°C for 15 min. Then they were moved to room temperature for another 15 min. After the recovery, slices were sequentially transferred to ACSF at room temperature containing either vehicle (DMSO) or 7,8-DHF for 1.5h (2h treatments) or 3.5h (4h treatments). Whole-cell patch-clamp measurements of E_GABA_ were performed under Cl^−^ load, using borosilicate patch pipettes with a resistance of 3 to 4 MΩ when filled with an intrapipette solution containing (in mM): 115 K-methylsulfate, 25 KCl, 2 MgCl_2_, 10 HEPES, 4 ATP-Na, 0.4 GTP-Na and 0.1% Lucifer Yellow (dipotassium salt) (pH adjusted to 7.2 with KOH). Data were low-pass filtered during acquisition at 3 kHz and sampled at 10 kHz using pClamp 10 (Molecular Devices). All recordings were performed at room temperature in standard ACSF supplemented with 10 µM CNQX, 40 µM APV, 1µM strychnine and 1 µM TTX. In experiments with 7,8-DHF-treated slices, the ACSF was supplemented with 10 µM 7,8-DHF, otherwise the ACSF was supplemented with equal concentration of DMSO. The slices were not used more than 1h after placing them in the recording solution. Only cells from lamina II of the spinal cord were targeted. Muscimol (500 µM) dissolved in HEPES-buffered ACSF, containing (in mM) 126 NaCl, 10 glucose, 2.5 KCl, 2 CaCl2, 2 MgCl2, 10mM HEPES (pH adjusted to 7.4), was pressure-applied (10psi) with a patch pipette using a picospritzer® III for 30 ms at different holding potentials. The tip of the pipette was placed approximately 5 μm from the center of the soma of the recording neurons. Only neurons with stable access resistance (lower than 22MΩ) and E_GABA_ during at least 2 recordings were included for subsequent analysis. Data analysis was performed offline with Clampfit 10.2 (Molecular Devices) and data was low-pass filtered at 1kHz. Experimental *E_GABA_* was extrapolated from the X-axis intersection of a linear regression (GraphPad Prism) from the I-V relationships of the GABA_A_ response to muscimol application. Measurements were adjusted to account for liquid junction potential (−8 mV).

### Cell surface biotinylation

The procedure for cell surface biotinylation assay in slices was conducted as previously described and validated ^7^ (Fig 1e and 5a). Briefly, after treatments, primary neurons or slices were washed two times in ice-cold PBS and then, incubated (1h; 4°C) with 1.5 mg/ml EZ-Link Sulfo-NHS-LC-Biotin (Thermo Scientific) in ice-cold PBS buffer, washed with ice-cold quenching solution (100 mM glycine in PBS) and lysed in 0.5 ml RIPA buffer (50 mM Tris-HCl pH 7.4, 150 mM NaCl, 1% Triton X-100, 0.5% Na-deoxycholate, and 0.1% SDS) supplemented with Complete® protease and PhosStop® phosphatase inhibitor cocktails (Roche). After incubation on a rotator (overnight; 4°C) and centrifugation at 18,000g (15 min; 4°C), the protein fraction was transferred into new tubes. After determination of total lysate protein concentration using the Bio-Rad DC Protein Assay, biotinylated cell surface proteins were purified from 500 µg total lysate proteins by adding 50 μl of slurry streptavidin agarose beads (Thermo Scientific) previously conditioned to RIPA buffer solution, incubating overnight at 4°C, and recovering the beads by centrifugation at 9,000g (1 min; 4°C). Finally, biotinylated cell surface proteins were released from the beads by boiling in 40 µl of 2x protein sample buffer and subjected to SDS-PAGE and Western blot analyses. β-actin, a non-plasmalemmal protein, was not detected in the biotinylated fraction, confirming the efficacy of the protocol (Supplemental Fig 1) ^7^. Protein fraction from the total lysates were used for total protein level assessments.

### SDS-PAGE and Western blot analyses

Total proteins (3-10 µg) and cell surface biotinylated-proteins (10-30 µl) were run on 7.5% TGX Stain-Free FastCast gels (Bio-Rad) and transferred onto PVDF membranes (Trans-Blot Turbo Transfer Pack; Bio-Rad) using the Trans-Blot Turbo Transfer System (Bio-Rad) for Western blot analyses. Before transferring, TGX Stain-Free gels were illuminated under UV (Biorad Gel Doc EZ Imager) to get Stain-Free images as input controls of the total lysate proteins (TLP). Blots were then sequentially incubated with primary and secondary antibodies and revealed using SuperSignal West Femto Maximum Sensitivity Substrate (Thermo Scientific) or Amersham ECL Western Blotting Detection Reagents (GE Healthcare). The next primary antibodies were applied overnight at 4°C: polyclonal rabbit anti-KCC2 (Millipore, #07-432) at a dilution of 1:1000 for membrane KCC2 and of 1:2000 for total KCC2; polyclonal rabbit anti-PARP (Cell Signaling, #9542) at a dilution of 1:500; monoclonal mouse anti-spectrin (Millipore, #MAB1622) at a dilution of 1:250; monoclonal rabbit anti-β-actin (Cell Signaling, #4970) at a dilution 1:10000 for the membrane protein negative control. The secondary antibodies were applied for 1h: Stabilized Peroxidase Conjugated Goat Anti-Rabbit (H+L) (Thermo Scientific, #32460); Stabilized Peroxidase Conjugated Goat Anti-Mouse (H+L) (Thermo Scientific, #31430) and Streptavidin-HRP (Cell Signaling Technologies, #3999S) were used at a dilution of 1:2000, 1:2000 and 1:3000, respectively.

### Immunoblot quantification

The mean intensity of immunoreactive bands on Western blots was measured using ImageJ software (NIH). Surface expression of KCC2 was evaluated by dividing biotinylated KCC2 proteins by total membrane biotinylated proteins (TMBP, Streptavidin-HRP images). Total expression of KCC2 was evaluated by dividing total KCC2 proteins by total lysate proteins (TLP, Strain-Free images). Since slices are cut in two pieces (i.e. internal control one and treated one), we normalized data a second time to express results relative to control by dividing surface and total ratios from treated samples by the ratio from their controls on the same gel. All Western blots from a given animal were conducted on the same gel.

### Ca2+ imaging in rat hippocampal neuronal cultures

Rat hippocampal primary cultures, on poly-d-lysine coated 12 mm coverslip (See *Primary neurons* methods section above), were transduced at 14 DIV with AAV2/PHP.eB-CAG-Twitch-2b ^49^(60000 viral genome copies per cell) (COVF Viral Vector Core (RRID:SCR_016477), https://tools.neurophotonics.ca/)/. The neurons were imaged after 19-23 DIV using a multiphoton videorate microscope (Bliq Photonique) with excitation wavelength 800 nm. Laser intensity was kept constant throughout the sets of experiments. Images were acquired at 31 images seconds using a water dipping Nikon 25x objective NA = 1.10. We collected the blue emission channel with ET480/40 (Chroma) and the green emission channel using ET540/40 (Chroma). A dichroic mirror FF509-di01 (Semrock) was used for splitting the signal in the two channels. To increase the signal, 31 consecutive images (1s) were averaged together. Cell regions of interest (ROIs) were chosen by the user and intensity ratios (ET540/40 channel over ET480/40 channel) were obtained by averaging the intensity values in the ROIs in the different images. For each image, a background ROI was delineated, and mean background values were subtracted in each individual channel.

Cultures treated with 7,8-DHF (10 μM) were preincubated in cultured media and observed at 1h, 2h, 4h post-incubation in ACSF. Treatments with NMDA (10 and 60 μM) were applied acutely, and neurons were incubated with NMDA on the microscope in circulating ACSF. Then, neurons were imaged after 15 min and 30 min of incubation.

### Rb+ influx assays

Defolliculated stage V–VI *Xenopus laevis* oocytes were micro-injected with 10 ng and 0,05 ng of cRNA coding for mouse KCC2b and mouse TrkB receptor, respectively, and maintained for 3 days at 18°C in Barth’s medium, supplemented with 125 μM furosemide and 200 nM k252a. Oocytes were preincubated for 15 min in Barth’s medium ^50^ containing either 50 or 200 µM 7,8-DHF or DMSO and the specific treatment (50 µM Dantrolene, 40 µM APV or 1 µM okadaic acid) and then incubated for another 45 min in a regular flux medium (in mM: 7 mM Rb^+^, 100 µM ouabain ^50^) containing the same treatment. At the end of the influx assay, oocytes were bathed three times in a wash solution supplemented with 250 µM furosemide and lysed in 25 µl pure nitric acid (one oocyte/vial). After complete drying (∼2h) at 80°C, 1 ml MilliQ water (Millipore) was added to each vial and Rb^+^ content was measured by atomic absorption spectrometry (spectrometer 240Z AA; Agilent Technologies). All steps were carried out at room temperature. Results are expressed as percentage change of Rb^+^ influx in compound-treated oocytes versus Rb^+^ influx in vehicle-treated oocytes. A variable slope [inhibitor] *vs*. response model with to constrain equal to 1 was fit using GraphPad Prism to the dose-response curve.

### Statistical analyses

All data are expressed as the mean ± S.E.M. unless otherwise stated. Parametric statistical analysis was used only when normality of the distribution was confirmed by D’Agostino & Pearson test. For non-parametric analysis, Wilcoxon signed-ranked tests were used for pairwise comparisons for single treatments and Kruskal-Wallis tests with post hoc Dunn’s test were used for comparisons between multiple independent treatments. For parametric analysis, student t-tests were used for comparisons from two independent treatments and analysis of variance (ANOVA) with post hoc Tukey’s tests were used for comparisons between multiple independent treatments. Sample sizes are consistent with those reported in similar studies. Values were considered outliers when detected by a ROUT’s test with a Q value of 0.1%. Differences were considered significant at *p* < 0.05. All the statistics were performed using GraphPad Prism.

## SUPPLEMENTAL INFORMATION

**Supplemental figure S1:**
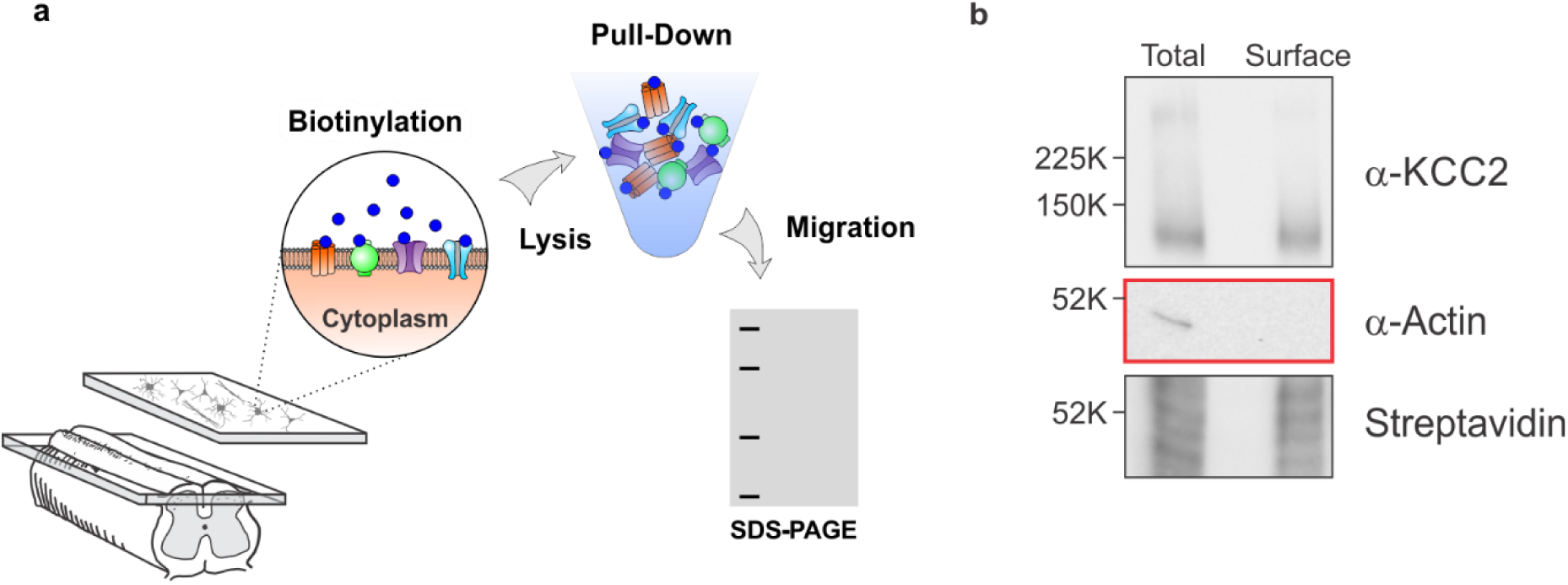
Validation of the biotinylation protocol used for membrane protein measurements on SDS-PAGE. (**a**) Schematic representation of the protocol used to purify plasmalemmal proteins in rat SDH slices. (**b**) Total and membrane fraction western blot results of a control sample. KCC2 (left) can be detected in both fractions. While β-actin (center), a cytosolic protein, can only be detected in the total fraction but it is absent in the membrane biotinylated protein fraction. The right panel shows the streptavidin input control used to detect the biotinylated protein in the sample.

**Supplemental figure S2:**
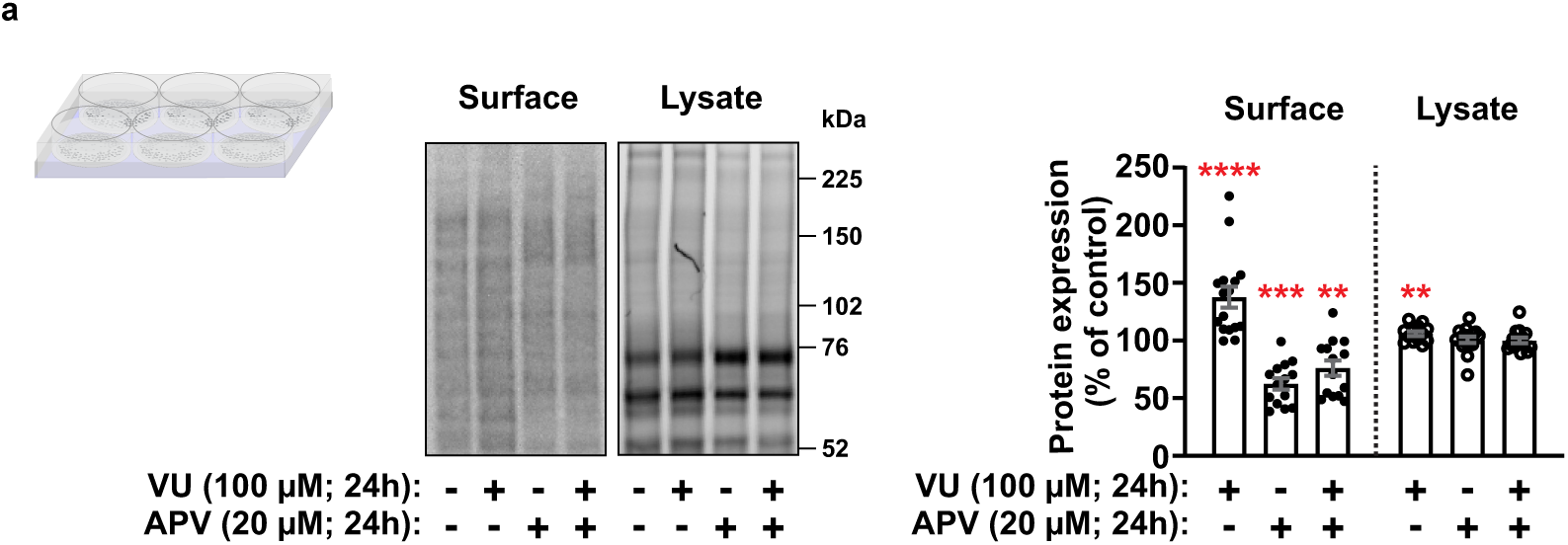
Input protein control observed in VU024551 and/or APV treated hippocampal cultures. **a**) Left panel: SDS-PAGE input images showing streptavidin in the surface samples, and stain free images for the total lysate fractions upon different treatments. Right panel: Quantification of the input controls in VU024551 and/or APV treated cultures. Significant changes in global membrane protein fraction can be observed upon the different treatments. Bars in the graphs represent the mean ± s.e.m. ** = p<0.01, *** = *p* < 0.001, **** = *p* < 0.0001.

## Notes

### Competing Interest Statement

The authors have declared no competing interest.

## REFERENCES

1. Doyon, N., Vinay, L., Prescott, S. A. & De Koninck, Y. Chloride Regulation: A Dynamic Equilibrium Crucial for Synaptic Inhibition. Neuron 89, 1157–1172 (2016).

2. Payne, J. A., Stevenson, T. J. & Donaldson, L. F. Molecular Characterization of a Putative K-Cl Cotransporter in Rat Brain: A NEURONAL-SPECIFIC ISOFORM. Journal of Biological Chemistry 271, 16245–16252 (1996).

3. Doyon, N. et al. Efficacy of Synaptic Inhibition Depends on Multiple, Dynamically Interacting Mechanisms Implicated in Chloride Homeostasis. PLOS Computational Biology 7, e1002149 (2011).

4. Doyon, N., Prescott, S. A. & De Koninck, Y. Mild KCC2 Hypofunction Causes Inconspicuous Chloride Dysregulation that Degrades Neural Coding. Frontiers in Cellular Neuroscience 9, (2016).

5. Boulenguez, P. et al. Down-regulation of the potassium-chloride cotransporter KCC2 contributes to spasticity after spinal cord injury. Nature Medicine 16, 302 (2010).

6. Ferrini, F. et al. Morphine hyperalgesia gated through microglia-mediated disruption of neuronal Cl− homeostasis. Nature Neuroscience 16, 183 (2013).

7. Gagnon, M. et al. Chloride extrusion enhancers as novel therapeutics for neurological diseases. Nat. Med. 19, 1524–1528 (2013).

8. Pathak, H. R. et al. Disrupted dentate granule cell chloride regulation enhances synaptic excitability during development of temporal lobe epilepsy. The Journal of neuroscience 27, 14012–14022 (2007).

9. Sullivan, C. R., Funk, A. J., Shan, D., Haroutunian, V. & McCullumsmith, R. E. Decreased Chloride Channel Expression in the Dorsolateral Prefrontal Cortex in Schizophrenia. PLOS ONE 10, e0123158 (2015).

10. Lorenzo, L.-E. et al. Enhancing neuronal chloride extrusion rescues α2/α3 GABAA-mediated analgesia in neuropathic pain. Nat. Commun. 11, 869 (2020).

11. Keramidis, I. et al. Restoring neuronal chloride extrusion reverses cognitive decline linked to Alzheimer’s disease mutations. Brain 146, 4903–4915 (2023).

12. Rivera, C. et al. BDNF-induced TrkB activation down-regulates the K+-Cl-cotransporter KCC2 and impairs neuronal Cl- extrusion. The Journal of cell biology 159, 747–752 (2002).

13. Zhang, Z., Wang, X., Wang, W., Lu, Y.-G. & Pan, Z. Z. Brain-derived neurotrophic factor-mediated downregulation of brainstem K+-Cl- cotransporter and cell-type-specific GABA impairment for activation of descending pain facilitation. Molecular pharmacology 84, 511–520 (2013).

14. Rivera, C. et al. Mechanism of Activity-Dependent Downregulation of the Neuron-Specific K-Cl Cotransporter KCC2. The Journal of Neuroscience 24, 4683–4691 (2004).

15. Coull, J. A. M. et al. BDNF from microglia causes the shift in neuronal anion gradient underlying neuropathic pain. Nature 438, 1017–1021 (2005).

16. Balkowiec, A. & Katz, D. M. Activity-Dependent Release of Endogenous Brain-Derived Neurotrophic Factor from Primary Sensory Neurons Detected by ELISAIn Situ. The Journal of Neuroscience 20, 7417–7423 (2000).

17. Balkowiec, A. & Katz, D. M. Cellular Mechanisms Regulating Activity-Dependent Release of Native Brain-Derived Neurotrophic Factor from Hippocampal Neurons. The Journal of Neuroscience 22, 10399–10407 (2002).

18. Aicardi, G. et al. Induction of long-term potentiation and depression is reflected by corresponding changes in secretion of endogenous brain-derived neurotrophic factor. Proceedings of the National Academy of Sciences of the United States of America 101, 15788–15792 (2004).

19. Bergami, M. et al. Uptake and recycling of pro-BDNF for transmitter-induced secretion by cortical astrocytes. The Journal of Cell Biology 183, 213–221 (2008).

20. Ji, Y. et al. Acute and gradual increases in BDNF concentration elicit distinct signaling and functions in neurons. Nature Neuroscience 13, 302–309 (2010).

21. Small, D. L. et al. Brain derived neurotrophic factor induction of N-methyl-D-aspartate receptor subunit NR2A expression in cultured rat cortical neurons. Neuroscience Letters 252, 211–214 (1998).

22. Leal, G., Comprido, D. & Duarte, C. B. BDNF-induced local protein synthesis and synaptic plasticity. Neuropharmacology 76, 639–656 (2014).

23. Puskarjov, M., Ahmad, F., Kaila, K. & Blaesse, P. Activity-Dependent Cleavage of the K-Cl Cotransporter KCC2 Mediated by Calcium-Activated Protease Calpain. The Journal of Neuroscience 32, 11356–11364 (2012).

24. Lee, H. H. C., Deeb, T. Z., Walker, J. A., Davies, P. A. & Moss, S. J. NMDA receptor activity downregulates KCC2 resulting in depolarizing GABAA receptor–mediated currents. Nature Neuroscience 14, 736 (2011).

25. Chamma, I. et al. Activity-Dependent Regulation of the K/Cl Transporter KCC2 Membrane Diffusion, Clustering, and Function in Hippocampal Neurons. The Journal of Neuroscience 33, 15488–15503 (2013).

26. Zhou, H.-Y. et al. N-methyl-D-aspartate receptor- and calpain-mediated proteolytic cleavage of K+-Cl- cotransporter-2 impairs spinal chloride homeostasis in neuropathic pain. The Journal of biological chemistry 287, 33853–33864 (2012).

27. Wang, W., Gong, N. & Xu, T.-L. Downregulation of KCC2 following LTP contributes to EPSP–spike potentiation in rat hippocampus. Biochemical and Biophysical Research Communications 343, 1209–1215 (2006).

28. Chorin, E. et al. Upregulation of KCC2 Activity by Zinc-Mediated Neurotransmission via the mZnR/GPR39 Receptor. The Journal of Neuroscience 31, 12916–12926 (2011).

29. Sakuragi, S., Tominaga-Yoshino, K. & Ogura, A. Involvement of TrkB-and p75(NTR)-signaling pathways in two contrasting forms of long-lasting synaptic plasticity. Scientific reports 3, 3185–3185 (2013).

30. Jang, S.-W. et al. A selective TrkB agonist with potent neurotrophic activities by 7,8-dihydroxyflavone. Proceedings of the National Academy of Sciences of the United States of America 107, 2687–2692 (2010).

31. Cazorla, M. et al. Identification of a low-molecular weight TrkB antagonist with anxiolytic and antidepressant activity in mice. J. Clin. Invest. 121, 1846–1857 (2011).

32. Lee, H. H. C. et al. Direct Protein Kinase C-dependent Phosphorylation Regulates the Cell Surface Stability and Activity of the Potassium Chloride Cotransporter KCC2. Journal of Biological Chemistry 282, 29777–29784 (2007).

33. Li, L. et al. Chloride Homeostasis Critically Regulates Synaptic NMDA Receptor Activity in Neuropathic Pain. Cell reports 15, 1376–1383 (2016).

34. Nguyen, H. T., Sawmiller, D. R., Wu, Q., Maleski, J. J. & Chen, M. Evidence supporting the role of calpain in the α-processing of amyloid-β precursor protein. Biochem. Biophys. Res. Commun. 420, 530–535 (2012).

35. Dutta, S., Chiu, Y. C., Probert, A. W. & Wang, K. K. W. Selective release of calpain produced αII-spectrin (α-fodrin) breakdown products by acute neuronal cell death. Biol. Chem. 383, 785–791 (2002).

36. Chaitanya, G. V., Steven, A. J. & Babu, P. P. PARP-1 cleavage fragments: signatures of cell-death proteases in neurodegeneration. Cell Commun. Signal. 8, 31 (2010).

37. Gilliams-Francis, K. L., Quaye, A. A. & Naegele, J. R. PARP cleavage, DNA fragmentation, and pyknosis during excitotoxin-induced neuronal death. Exp. Neurol. 184, 359–372 (2003).

38. Busciglio, J. & Yankner, B. A. Apoptosis and increased generation of reactive oxygen species in Down’s syndrome neurons in vitro. Nature 378, 776–779 (1995).

39. Dedek, A. et al. Loss of STEP61 couples disinhibition to N-methyl-d-aspartate receptor potentiation in rodent and human spinal pain processing. Brain 142, 1535–1546 (2019).

40. Hildebrand, M. E. et al. Potentiation of Synaptic GluN2B NMDAR Currents by Fyn Kinase Is Gated through BDNF-Mediated Disinhibition in Spinal Pain Processing. Cell Reports 17, 2753–2765 (2016).

41. Caldeira, M. V. et al. BDNF regulates the expression and traffic of NMDA receptors in cultured hippocampal neurons. Molecular and Cellular Neuroscience 35, 208–219 (2007).

42. Lin, S.-Y. et al. BDNF acutely increases tyrosine phosphorylation of the NMDA receptor subunit 2B in cortical and hippocampal postsynaptic densities. Molecular Brain Research 55, 20–27 (1998).

43. Yoshii, A. & Constantine-Paton, M. Postsynaptic BDNF-TrkB signaling in synapse maturation, plasticity, and disease. Developmental neurobiology 70, 304–322 (2010).

44. MacDermott, A. B., Mayer, M. L., Westbrook, G. L., Smith, S. J. & Barker, J. L. NMDA-receptor activation increases cytoplasmic calcium concentration in cultured spinal cord neurones. Nature 321, 519–522 (1986).

45. Díaz, D., Bartolo, R., Delgadillo, D. M., Higueldo, F. & Gomora, J. C. Contrasting Effects of Cd2+ and Co2+ on the Blocking/Unblocking of Human Cav3 Channels. The Journal of Membrane Biology 207, 91–105 (2005).

46. Kasai, H. & Neher, E. Dihydropyridine-sensitive and omega-conotoxin-sensitive calcium channels in a mammalian neuroblastoma-glioma cell line. The Journal of Physiology 448, 161–188 (1992).

47. Wakamori, M. et al. Functional Characterization of Ion Permeation Pathway in the N-Type Ca2+ Channel. Journal of Neurophysiology 79, 622–634 (1998).

48. Thévenod, F. & Jones, S. W. Cadmium block of calcium current in frog sympathetic neurons. Biophysical Journal 63, 162–168 (1992).

49. Thestrup, T. et al. Optimized ratiometric calcium sensors for functional in vivo imaging of neurons and T lymphocytes. Nat. Methods 11, 175–182 (2014).

50. Bergeron, M. J., Gagnon, É., Caron, L. & Isenring, P. Identification of Key Functional Domains in the C Terminus of the K+-Cl– Cotransporters. Journal of Biological Chemistry 281, 15959–15969 (2006).

51. Rogalski, S. L., Appleyard, S. M., Pattillo, A., Terman, G. W. & Chavkin, C. TrkB Activation by Brain-derived Neurotrophic Factor Inhibits the G Protein-gated Inward Rectifier Kir3 by Tyrosine Phosphorylation of the Channel. Journal of Biological Chemistry 275, 25082–25088 (2000).

52. Ferrini, F., Lorenzo, L.-E., Godin, A. G., Quang, M. L. & De Koninck, Y. Enhancing KCC2 function counteracts morphine-induced hyperalgesia. Scientific Reports 7, 3870 (2017).

53. Silayeva, L. et al. KCC2 activity is critical in limiting the onset and severity of status epilepticus. Proceedings of the National Academy of Sciences 112, 3523–3528 (2015).

54. Tillman, L. & Zhang, J. Crossing the Chloride Channel: The Current and Potential Therapeutic Value of the Neuronal K+-Cl- Cotransporter KCC2. BioMed Research International 2019, 12 (2019).

55. Schulte, J. T., Wierenga, C. J. & Bruining, H. Chloride transporters and GABA polarity in developmental, neurological and psychiatric conditions. Neuroscience & Biobehavioral Reviews 90, 260–271 (2018).

56. Bos, R. et al. Activation of 5-HT2A receptors upregulates the function of the neuronal K-Cl cotransporter KCC2. Proceedings of the National Academy of Sciences 110, 348–353 (2013).

57. Weber, M., Hartmann, A.-M., Beyer, T., Ripperger, A. & Nothwang, H. G. A Novel Regulatory Locus of Phosphorylation in the C Terminus of the Potassium Chloride Cotransporter KCC2 That Interferes with N-Ethylmaleimide or Staurosporine-mediated Activation. Journal of Biological Chemistry 289, 18668–18679 (2014).

58. Moore, Y. E., Deeb, T. Z., Chadchankar, H., Brandon, N. J. & Moss, S. J. Potentiating KCC2 activity is sufficient to limit the onset and severity of seizures. Proceedings of the National Academy of Sciences 115, 10166–10171 (2018).

59. Vale, C., Schoorlemmer, J. & Sanes, D. H. Deafness Disrupts Chloride Transporter Function and Inhibitory Synaptic Transmission. The Journal of Neuroscience 23, 7516–7524 (2003).

60. Chaplan, S. R., Malmberg, A. B. & Yaksh, T. L. Efficacy of Spinal NMDA Receptor Antagonism in Formalin Hyperalgesia and Nerve Injury Evoked Allodynia in the Rat. Journal of Pharmacology and Experimental Therapeutics 280, 829–838 (1997).

61. Coull, J. A. M. et al. Trans-synaptic shift in anion gradient in spinal lamina I neurons as a mechanism of neuropathic pain. Nature 424, 938 (2003).

62. Chamma, I., Chevy, Q., Poncer, J. C. & Levi, S. Role of the neuronal K-Cl co-transporter KCC2 in inhibitory and excitatory neurotransmission. Front. Cell. Neurosci. 6, (2012).

63. Barthó, P., Payne, J. A., Freund, T. F. & Acsády, L. Differential distribution of the KCl cotransporter KCC2 in thalamic relay and reticular nuclei. Eur. J. Neurosci. 20, 965–975 (2004).

64. Al Awabdh, S. et al. Gephyrin interacts with the K-Cl cotransporter KCC2 to regulate its surface expression and function in cortical neurons. J. Neurosci. 42, 166–182 (2022).

65. Heubl, M. et al. GABAA receptor dependent synaptic inhibition rapidly tunes KCC2 activity via the Cl--sensitive WNK1 kinase. Nat. Commun. 8, 1776 (2017).

66. Donneger, F., et al. Enhancing KCC2 function reduces interictal activity and prevents seizures in temporal lobe epilepsy. bioRxiv (2023) doi:10.1101/2023.09.16.557753.

67. Perez-Sanchez, J. & De Koninck, Y. Regulation of Chloride Gradients and Neural Plasticity. Oxford Research Encyclopedia of Neuroscience (2019) doi:10.1093/acrefore/9780190264086.013.238.

68. Hewitt, S. A., Wamsteeker, J. I., Kurz, E. U. & Bains, J. S. Altered chloride homeostasis removes synaptic inhibitory constraint of the stress axis. Nature Neuroscience 12, 438–443 (2009).

69. Grau, J. W. & Huang, Y.-J. Metaplasticity within the spinal cord: Evidence brain-derived neurotrophic factor (BDNF), tumor necrosis factor (TNF), and alterations in GABA function (ionic plasticity) modulate pain and the capacity to learn. Neurobiology of Learning and Memory 154, 121–135 (2018).

70. Nault, F. & De Koninck, P. Dissociated Hippocampal Cultures. in Protocols for Neural Cell Culture: Fourth Edition (ed. Doering, L. C.) 137–159 (Humana Press, Totowa, NJ, 2009).

71. Rivera, C. et al. The K+/Cl− co-transporter KCC2 renders GABA hyperpolarizing during neuronal maturation. Nature 397, 251 (1999).

72. Chéry, N., Yu, X. H. & De Koninck, Y. Visualization of lamina I of the dorsal horn in live adult rat spinal cord slices. Journal of Neuroscience Methods 96, 133–142 (2000).

73. Digman, M. A., Caiolfa, V. R., Zamai, M. & Gratton, E. The Phasor Approach to Fluorescence Lifetime Imaging Analysis. Biophysical Journal 94, L14–L16 (2008).

